# Determining the architecture of nuclear ring of *Xenopus laevis* nuclear pore complex using integrated approaches

**DOI:** 10.1101/2021.11.10.468004

**Authors:** He Ren, Linhua Tai, Yun Zhu, Chun Chan, Qun Zhao, Jiashu Xu, Xiangyang Wang, Xinyao Ma, Lili Zhao, Xiaojun Huang, Guoliang Yin, Mingkang Jia, Xiaohong Zhu, Yuxin An, Gan Zhao, Changxiong Huang, Lihua Zhang, Jun Fan, Fei Sun, Chuanmao Zhang

## Abstract

The nuclear pore complexes (NPCs) are large protein assemblies as a physical gate to regulate nucleocytoplasmic transport. Here, using integrated approaches including cryo-electron microscopy, hybrid homology modeling and cell experiment, we determined the architecture of the nuclear ring (NR) from *Xenopus laevis* oocytes NPC at subnanometer resolution. In addition to the improvement of the Y complex model, eight copies of Nup205 and ELYS were assigned in NR. Nup205 connects the inner and outer Y complexes and contributes to the assembly and stability of the NR. By interacting with both the inner Nup160 and the nuclear envelope (NE), the N-terminal β-propeller and α-solenoid domains of ELYS were found to be essential for accurate assembly of the NPC on the NE.

## INTRODUCTION

Nuclear pore complexes (NPCs) are large protein assemblies constructed from multiple copies of approximately 30 proteins that span the double-layered nuclear envelope (NE) and form gates controlling the exchange of macromolecules between the nucleus and the cytoplasm ^1^. A fully assembled NPC contains approximately 550 protein subunits termed nucleoporins (Nups) in fungi and approximately 1,000 Nups in vertebrates ^2,3^. The molecular mass of the NPC is approximately 52 MDa in yeast and 125 MDa in higher eukaryotes, making the NPC one of the largest biomacromolecular assemblies in eukaryotic cells ^3–5^. The number of NPCs within one cell varies greatly among different species, from 200 per nucleus in yeast to 2,000-5,000 per somatic cell nucleus in vertebrates to ~5 × 10^7^ copies per oocyte nucleus in *Xenopus laevis* (*X. laevis*) ^4^.

NPCs were first discovered in the early 1950s when electron microscopy was applied to examine amphibian oocyte nuclei ^6^. In recent decades, both cryo-electron tomography (cryo-ET) in conjunction with subtomogram averaging (STA) and X-ray crystallography have been applied to determine the molecular structures of NPCs. Morphologically, a fully assembled NPC exhibits eightfold rotational symmetry along the axis perpendicular to the NE, and each of the asymmetric units contains nucleoplasmic and cytoplasmic subunits, which are joined at the equator of the NPC ^4^. The main scaffold of a mature NPC includes a cytoplasmic ring (CR), an inner ring (IR), a luminal ring (LR) and a nuclear ring (NR), as well as other functional domains, including cytoplasmic filaments, a permeability barrier formed by phenylalanine-glycine (FG) repeat-rich Nups, and the nuclear basket ^2,7–10^. Based on the structural analysis of NPC components^11–16^ and Y-complexes^7,17,18^, the overall structures of the NPCs in several species, including *Saccharomyces cerevisiae* (*S. cerevisiae*), *Schizosaccharomyces pombe* (*S. pombe*), *Dictyostelium discoideum* (*D. discoideum*), *Chlamydomonas reinharadtii* (*C. reinharadtii*), *X. laevis* and *Homo sapiens* (*H. sapiens*), have been studied at nanometer resolution via cryo-ET along with STA ^3,8,9,11,12,19–22^ and revealed that the backbones of the CR and NR are formed by the Y-shaped Nup84 complexes in fungi and the Nup107 complexes in vertebrates^5,23^.

More recently, the structure of the CR in *X. laevis* oocyte NPCs was resolved at resolutions of 5.5-7.9 Å on average by a cryo-electron microscopy (cryo-EM) single particle analysis (SPA) approach, providing structural details and assigning additional Nups (Nup205, Nup214 complex and Nup358 complex) in addition to the Y-shaped Nup107 complexes in CR ^21^. However, to reveal the structural differences between the CR and NR and understand how ELYS (embryonic large molecule derived from yolk sac, also known as Mel-28 or AHCTF1), an essential Nup for postmitotic NPC assembly ^9,24^, specifically localizes to the NR, as well as to determine how the NR interacts with the nuclear basket, a higher resolution structure of the NR needs to be obtained.

In this work, after revisiting the structures of the NPC components from *X. laevis* oocytes via both cryo-ET STA and cryo-EM SPA approaches, we focused on analyzing the detailed structure of the NR by further utilizing hybrid homology modeling approaches. The overall resolution of the resulting NR structure is 7.8 Å with the core region at 6.8 Å resolution, which enabled us to build the more complete model of the NR. 16 Y-shaped Nup107 subcomplexes forming inner and outer rings in NR, while only 8 copies of Nup205 were observed in the NR. We identified and modeled ELYS, which interacts with the NE via its N-terminal β-propeller domain and attaches to the inner Nup160 only via its α-solenoid domain. In addition, we observed a region of unassigned densities that might represent the locations of portions of TPR and Nup153, the components of the nuclear basket. Overall, our work provides an advance in understanding the architecture, assembly, and function of the NPCs.

## RESULTS

### Overall structure of the *X. laevis* oocyte NPC

To maintain the integrity of the NPC structure, we performed a structural study of intact NPCs on NEs directly using *X. laevis* oocyte nuclei. The NEs were manually isolated from stage VI oocyte nuclei of *X. laevis*. The isolation and NE preparation procedures were verified by in-lens field emission scanning electron microscopy (In-Lens FESEM) (Fig. 1A–B) and resin-embedded transmission electron microscopy (TEM) (Fig. 1C), which showed an integrated architecture with visible cytoplasmic filaments and well-ordered nuclear baskets (Fig. 1A–D) ^25,26^ We performed both cryo-EM SPA and cryo-ET STA to reconstruct the 3D structures of the NPCs and reached resolutions of 29 Å and 65 Å for scaffold rings, respectively (Fig. S1). Considering the flexibility and dynamics of the whole NPC, the resolution could not be improved further by global structural averaging. Thus, we applied a block-based cryo-EM SPA approach by masking different local regions, which was similar to previous reports ^21,27^, and successfully resolved the cryo-EM maps of asymmetric units of the CR, NR and IR at resolutions of 7.6 Å, 6.8-10.6 Å and 9.8 Å, respectively (Fig. 1 E–G, Fig. S2, Fig. S3 and Movie S1). In addition to the overall map of the NR asymmetric unit resolved at 7.8 Å, a further tight mask was applied to increase the resolution of the stable core region of NR to 6.8 Å. In addition, another mask focusing on the unknown density region (UDR) of the NR was applied to achieve a map at a resolution of 10.6 Å (Fig. S3).

**Fig. 1.**
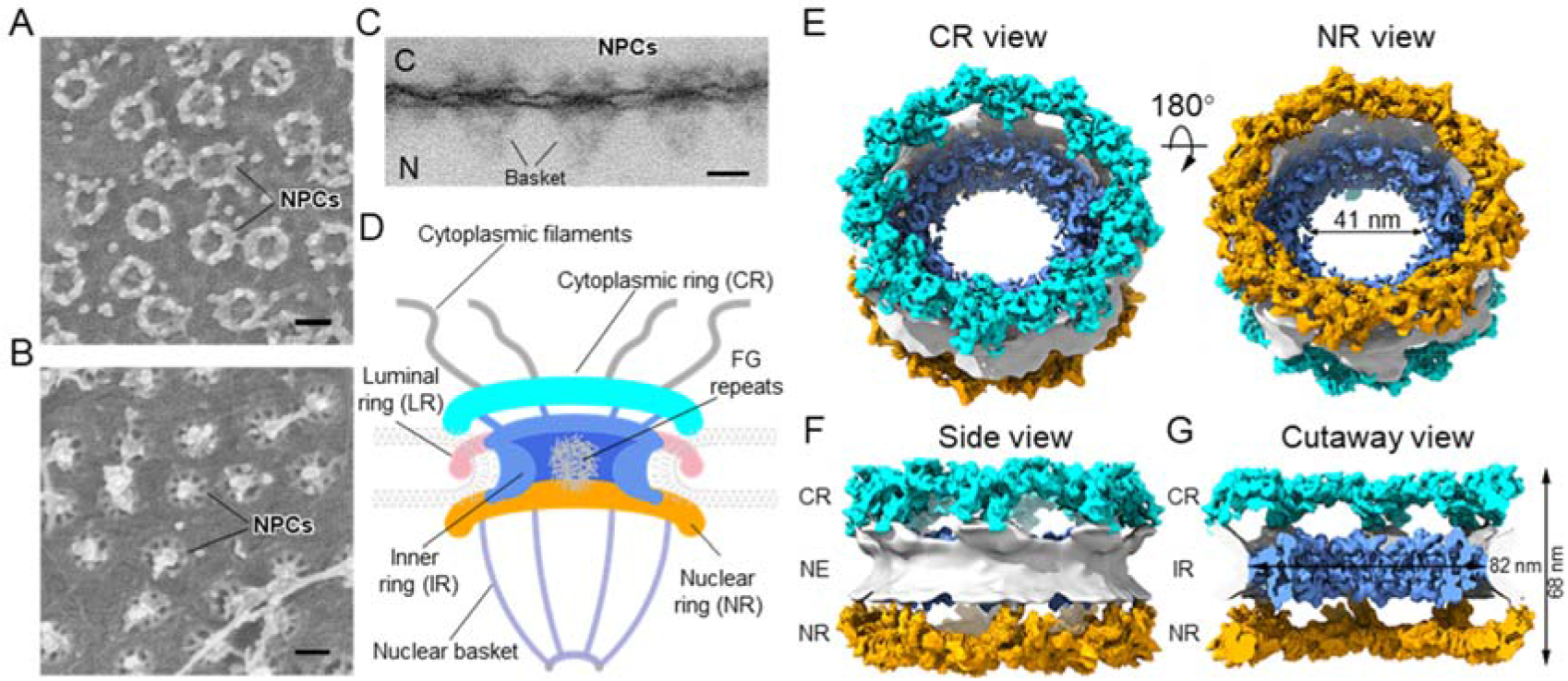
Structure of the *X. laevis* oocyte NPC NR. **(A)** High magnification of the cytoplasmic side surface of the NE *X. laevis* oocytes imaged by In-Lens FESEM. **(B)** Nucleoplasmic side surface of the *X. laevis* oocyte NE imaged by In-Lens FESEM. (**C)** TEM image of the resin-embedded *X. laevis* oocyte NE. The section was cut perpendicular to the NE. C indicates the cytoplasmic side, and N indicates the nucleoplasmic side. Scale bars, 100 nm in (**A-C). (D)** A cutaway schematic representation of a fully assembled NPC. The main components of the NPC include the cytoplasmic filaments, CR in cyan, LR in light pink, FG repeats in gray, IR in cornflower blue, NR in orange and nuclear basket in violet. (**E**) Overall views of the cryo-EM map of *X. laevis* NPCs from the CR (cyan) and NR (orange) sides. **(F & G)** Side and cutaway views of the *X. laevis* NPC, showing the IR in cornflower blue and NE in light gray.

Overall, the height of *X. laevis* NPCs from CR to NR was 68 nm, and the outer diameters of both the NR and CR were 125 nm, while the inner diameters of the NR and CR region were 75 and 68 nm, respectively. In addition, the outer/inner diameter of *X. laevis* NPCs in the IR region was measured as 82/41 nm (Fig. 1E–G). The overall shape of *X. laevis* NPCs is similar to that of *H. sapiens* NPCs reported from HeLa cells ^7^.

### Improved model of NR from *X. laevis* NPC

Based on the subnanometer resolution of the NR map, we built a more complete model of NR from *X. laevis* NPC (Fig. 2). It is worth noting that the map still has moderate anisotropic resolution due to the imperfect Fourier space sampling (Fig. S2 and Fig. S3) in data collection, but many secondary structural elements of NR components can be identified (Movie S1), similar as the CR study in previous report ^28^. In addition to all the subunits of the Y complex (Seh1, Sec13, Nup37, Nup43, Nup85, Nup96, Nup107, Nup133, and Nup160) that identified in CR, we also observed additional densities that were later assigned as Nup205 and ELYS, as well as an unknown density region (UDR) that sits on the nucleoplasmic side of the NR (Fig. 2D).

**Fig. 2.**
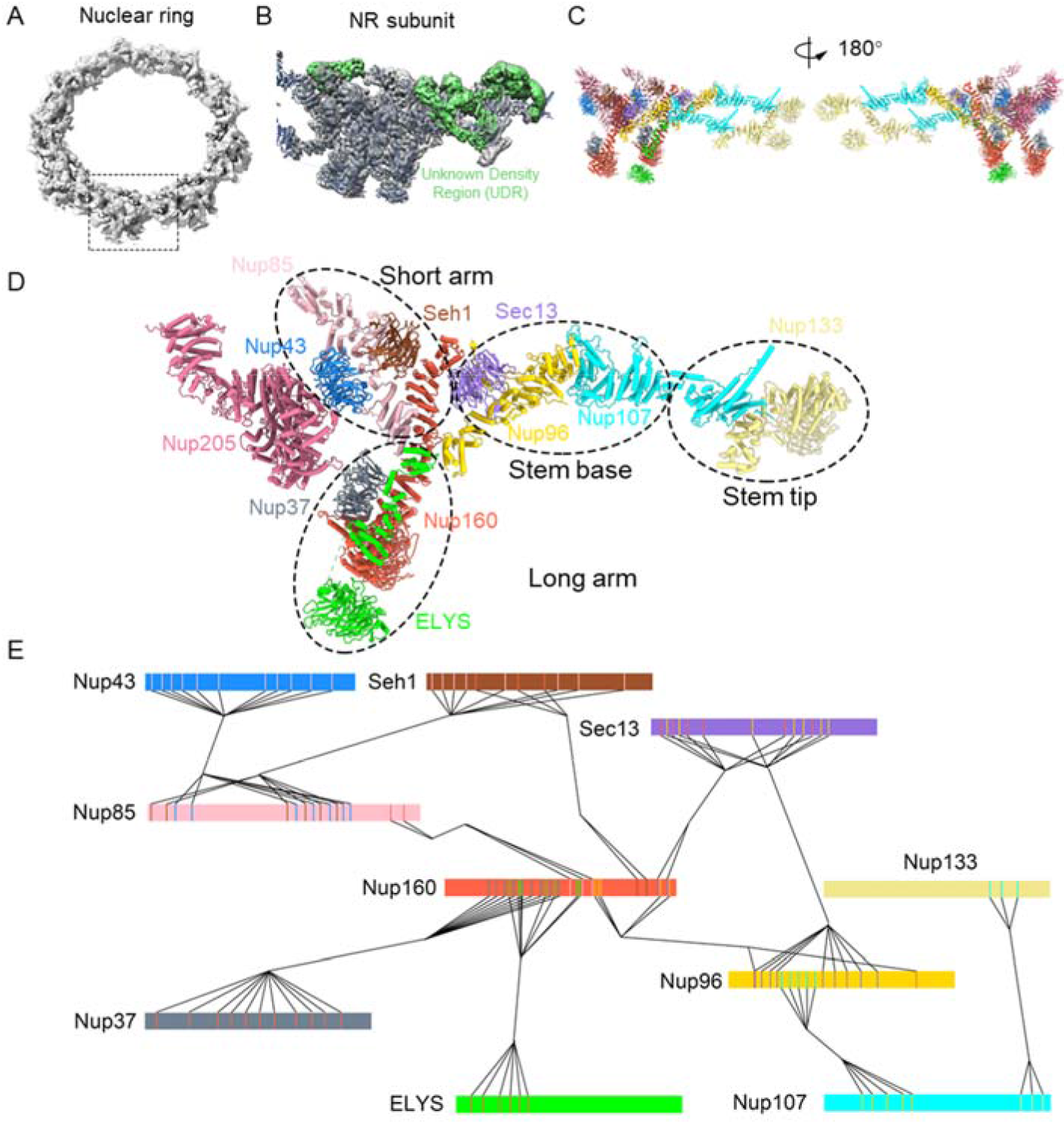
Model of the Y complex in *X. laevis* NR. **(A)** Overall reconstructed map of the NR with a tilting angle of 60°. **(B)** Cryo-EM map of the NR asymmetric unit with the fitted model and UDR shown in green. **(C)** Model of two Y complexes with Nup205 in different views. Various colors indicate different Nups. **(D)** The model of the inner Y complex and Nup205. The short arm comprises Nup85, Nup43 and Seh1, the long arm comprises Nup37, Nup160 and ELYS, the stem base comprises Sec13, Nup96 and part of Nup107, and the stem tip comprises part of Nup107 and Nup133. **(E)** Model of contact regions in the inner Y complex. The interactions between different Nups are indicated by black lines. The corresponding interaction sites are shown as vertical lines on each Nup accordingly.

Starting from the reported models of the *H. sapiens* Y complex (PDB code 5A9Q) ^9^, we managed to build a more complete model of the *X. laevis* Y complex (Fig. 2C) using map-based homology modeling and molecular dynamics flexible fitting (MDFF) approaches. To the best of our knowledge, this is the most complete model of the Y complex from previous reports. Similar to that of *H. sapiens* NPCs ^7,9^, the NR of *X. laevis* NPCs also contains 16 copies of Y complexes forming two layers of rings. Each eightfold asymmetric unit of the NR is made up of two copies of Y complexes, the inner and outer ones (Fig. 2C). Consistently, the Y complex comprises three parts: a short arm containing Nup85, Nup43 and Seh1, a long arm containing Nup160 and Nup37, and a stem containing Nup96, Sec13, Nup107 and Nup133 (Fig. 2D). Overall, we determined that the Y complex is mainly composed of five α-solenoid domains and six β-propeller domains in the NR of *X. laevis* NPC (Fig. 2D).

With our update model of the Y complex, we are able to analyze the detailed contact interfaces among different subunits (Fig. 2E), many of which have been reported in *H. sapiens* NPCs ^7,9^. In particular, with the completion and refinement of the C-terminal parts of Nup85, Nup160 and Nup96, their interaction regions were assigned. The C-terminal region of Nup85 forms contacts with the middle region of Nup160. The middle region and the C-terminal region of Nup160 make contacts with the N-terminal region and C-terminal region of Nup96. The interaction between Nup160 and Nup96 is stabilized by Sec13, which forms substantial contacts with the C-terminal region of Nup160 and the N-terminal and middle regions of Nup96 (Fig.2E).

### Comparison between the NR and the CR

The diversity of the Y complex has been described before ^9,21,29^. It would be of interest to investigate whether the CR and NR share the same conformation in *X. laevis* NPC. To keep consistency of scale, we compared the NR and CR based on our cryo-EM maps at 7.6 Å resolution for CR and 7.8 Å resolution for NR. To reveal the difference between the CR and NR, we computed their difference maps and superimposed them onto the update model of the NR. We found that there is no obvious density in the difference maps overlapping with the model. This observation suggests that the Y complexes in both NR and CR exhibit almost the same architecture, which agrees with previous results ^7,9^, and that our composite model of the NR can be directly docked into the density of the CR map with high confidence level (Fig. 3C–D).

**Fig. 3.**
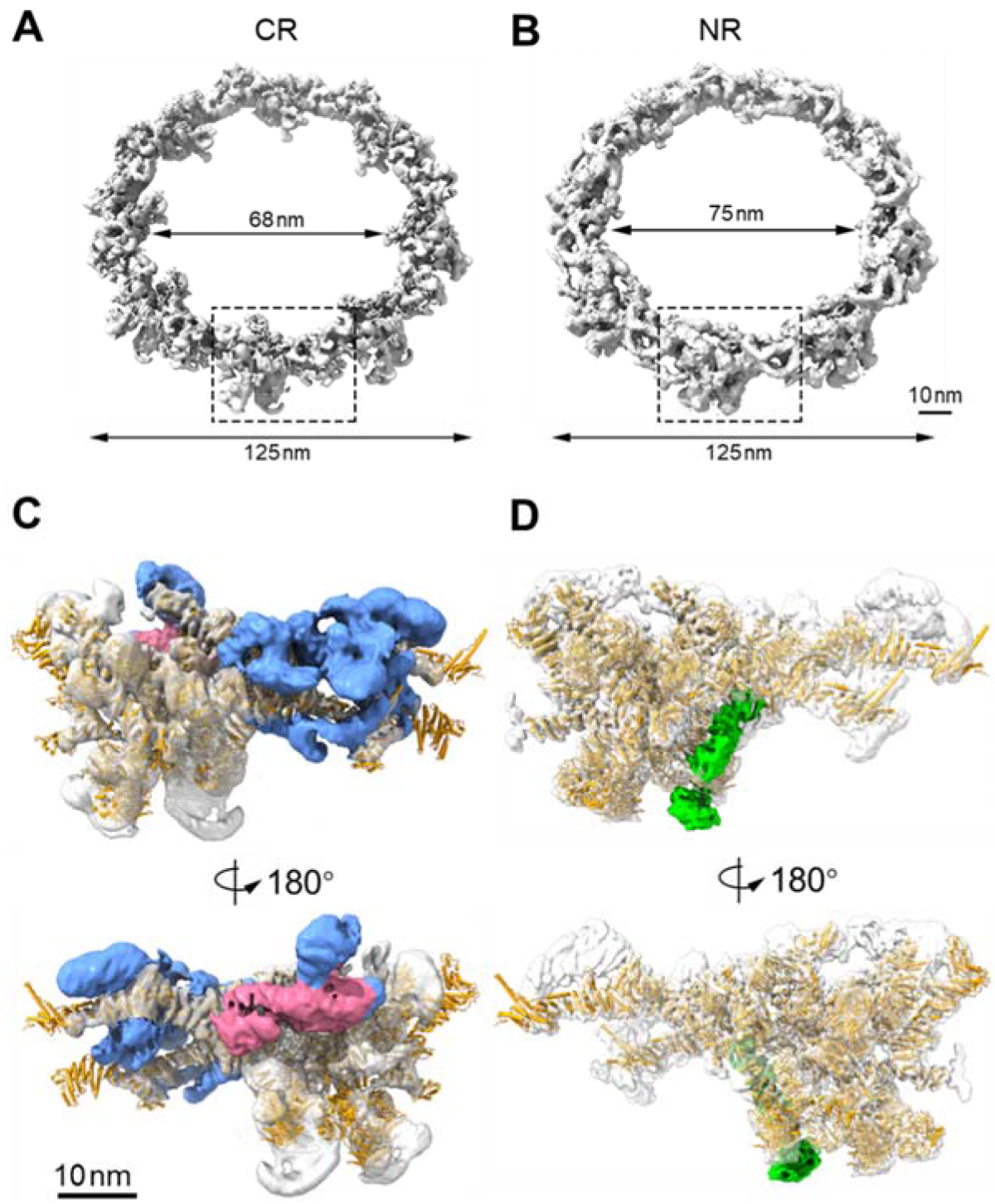
Comparing CR and NR asymmetric unit. (A & B) An overall reconstructed map of the *X. laevis* NPC CR and NR, respectively. The dashed rectangle indicates the asymmetric unit. The map of the CR asymmetric unit with the model of Y complexes (orange) fitted. The difference map obtained by subtracting the NR from the CR is shown in cornflower blue for the region of Nup358 and Nup214 complexes and in pale violet red for the region of inner Nup205. The map of the NR asymmetric unit with the model of Y complexes (orange) fitted. The difference map obtained by subtracting the CR from the NR is shown in green for the ELYS region.

In addition, our difference maps imply distinguished unique structures of both the CR and NR (Fig. 3C–D). On the difference map obtained by subtracting the NR from the CR in each asymmetric unit, we observed that the significant lump densities mainly wrap around the stem regions of both the outer and inner Y complexes, which correspond to the location of Nup358 and Nup214 complexes according to a recent report ^21^. In addition, we found another block of density difference located near the inner Nup85 and Nup160 (Fig. 3C) that is consistent with the structure assigned to inner Nup205 by a recent report ^21^. However, although it was reported that there are two copies of Nup205 in one asymmetric unit of the CR, that is, an inner copy and an outer copy ^21^, we observed only the inner one in our difference map, suggesting that there is only one copy of Nup205 in the NR and that this single Nup205 in the NR should occupy a position similar to that of its counterpart in the CR (see below).

On the difference map obtained by subtracting the CR from the NR in each asymmetric unit, there is an obvious β-propeller domain and adjacent α-solenoid domain situated by the side of the inner Nup160 near the NE (Fig. 3D). Considering that ELYS is an unique component of the NR that contains an N-terminal β-propeller domain and a connected α-solenoid domain and interacts with both Nup160 and the NE ^2^, we assigned this density difference to ELYS and built its model (see below).

### Nup205 and the NR assembly

In NR subunit map, we observed a clear density sandwiched between the inner and outer Y complexes in proximity to the short arm of the outer Y complex (Fig. 4A). The equivalent density in the CR was previously assigned to Nup188 or Nup205 due to its interaction with the Nup214 complex *in vitro* ^9^ but was more recently suggested to be Nup205 ^21^. Based on our high-resolution map, we further verified the assignment of this density by using integrative structural modeling. Utilizing a cryo-EM map-based homology modeling approach, we modeled and fitted Nup205 and Nup188 into this density (Fig. 4B). We observed that although Nup205 and Nup188 are similar in overall molecular shape and are mainly composed of α-helices, their topologies are different. In particular, the unique long helix of Nup205, also named the tower helix ^5^, fits well with the density of the NR subunit map. Due to the lack of this long helix in Nup188, the fitting of Nup188 into the density of the NR subunit map is not well matched in the specific tower helix region (Fig. 4B). Moreover, we also revealed that the overall cross correlation (CC) of Nup205 (0.48) in the map was higher than that of Nup188 (0.11) ^30^. Based on these results, we assigned Nup205 to the density sandwiched between the inner and outer Y complexes and refined the model of Nup205 with higher structural accuracy.

**Fig. 4.**
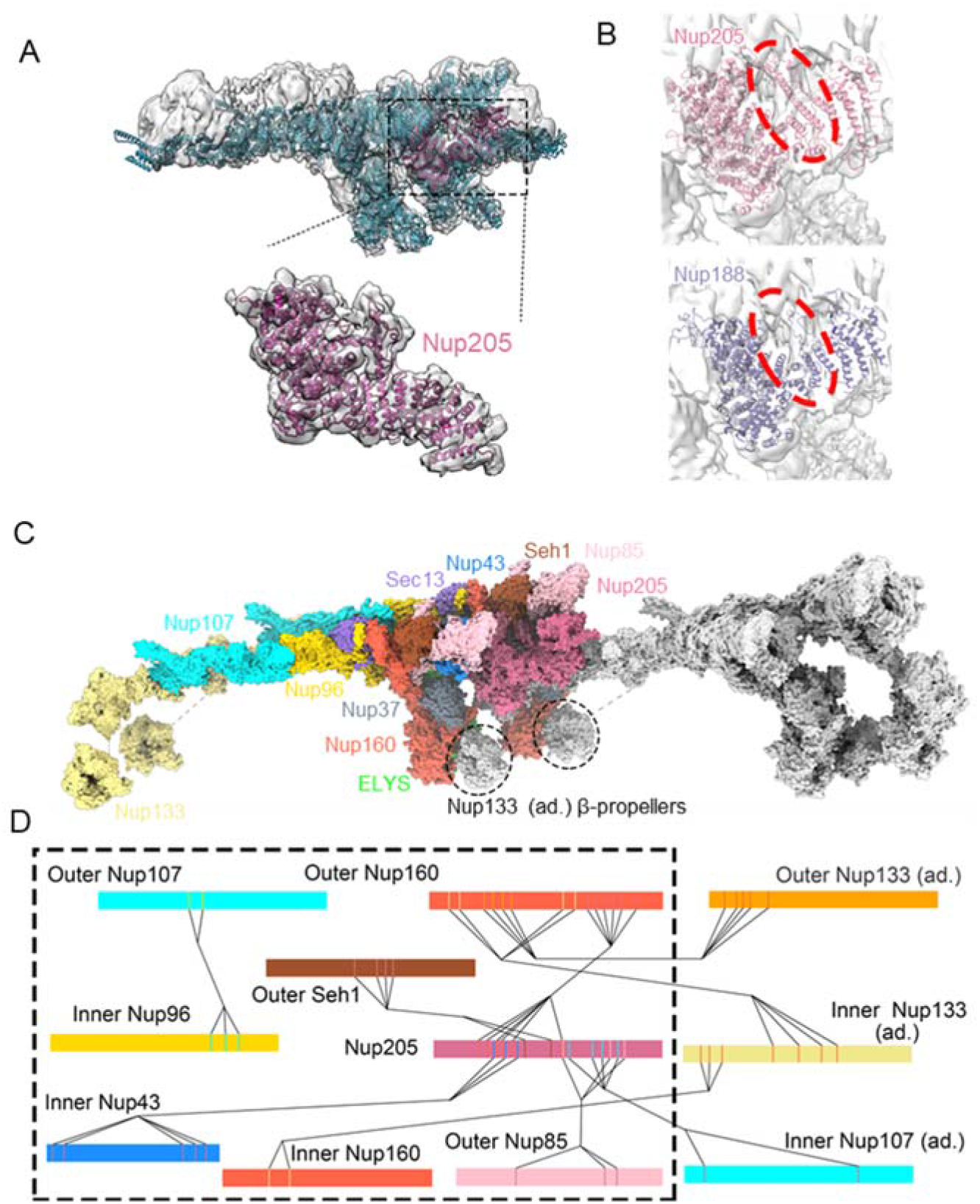
Identification of Nup205 and its interactions with NR components. **(A)** Identification and modeling of Nup205. The map of the NR asymmetric unit (gray) is fitted with the model of Y complexes and Nup205 (violet red). (**B**) Comparing the modeling of Nup205 (pale violet red) and Nup188 (dark slate blue) at the same region, indicating the Nup205 tower helix region. **(C)** An overall view of the Nup contact interfaces among the Y complexes within two NR asymmetric units. (**D**) Model of contact regions among different Y complexes. The interactions between different Nups are indicated by black lines. The corresponding interaction sites are shown as vertical lines on each Nup accordingly. The dashed rectangle indicates the Nups in one NR asymmetric unit.

With assignment of Nup205 to the specific region between the inner and the outer Y complexes, we investigated how the subcomplexes are connected with each other and revealed that the connections at the short and long arm regions are mediated by Nup205 (Fig. 4C–D). We discovered that while the N-terminal region of Nup205 has contacts with the N- and C-terminal regions of the inner Nup43, two domains in the middle region of Nup205 form an interaction interface with the middle region of the outer Seh1. A large portion of Nup205 from the N-terminal to the middle regions makes extensive contact with the C-terminal regions of the outer Nup160. The middle region and the C-terminal region of Nup205 contact with N-terminal and middle regions of the outer Nup85. With these connections, Nup205 plays a crucial role in assembling the two Y complexes into one asymmetric unit of the NR. In addition to the role of Nup205 as a hub, direct interactions also occur between the stem regions of the inner and outer Y complexes, where the middle regions of the outer Nup107 interact with the C-terminal regions of the inner Nup96. These interactions further stabilize the structure of the NR asymmetric units at their stem parts.

Next, we studied the assembly of the NR with the asymmetric units of Y complexes. Based on our subnanometer resolution cryo-EM map, we found out that the connection of adjacent asymmetric units are mediated by four major interaction pairs, including Nup205 and the adjacent inner Nup107, the outer Nup160 and the adjacent inner Nup133, the outer Nup160 and the adjacent outer Nup133, the inner Nup160 and the adjacent inner Nup133 (Fig. 4D). The Nup133 not only anchors the Y complexes onto the NE but also acts as a linker to assist Nup205 in the head-to-tail arrangement of Y complexes of the NR ^9,31,32^.

### Modeling of ELYS and its role in the assembly of the NPC

ELYS is a large chromatin-associated protein with an AT-hook DNA binding motif and is required for postmitotic NPC assembly ^24,33,34^. Depletion of ELYS leads to severe disruption of the NPC on the NE ^24,33–35^. ELYS roughly comprises an N-terminal β-propeller domain, a middle α-solenoid domain and a C-terminal unstructured region (Fig. 5A) ^2^. The crystal structure of the β-propeller domain of ELYS has been previously resolved at 1.9 Å resolution ^36^. It has been suggested that vertebrate ELYS is located in the region of the NR and interacts with Nup160 ^9,24^. However, the exact location and orientation of ELYS’ α-solenoid domain on the NR and its full-length structure remain elusive.

**Fig. 5.**
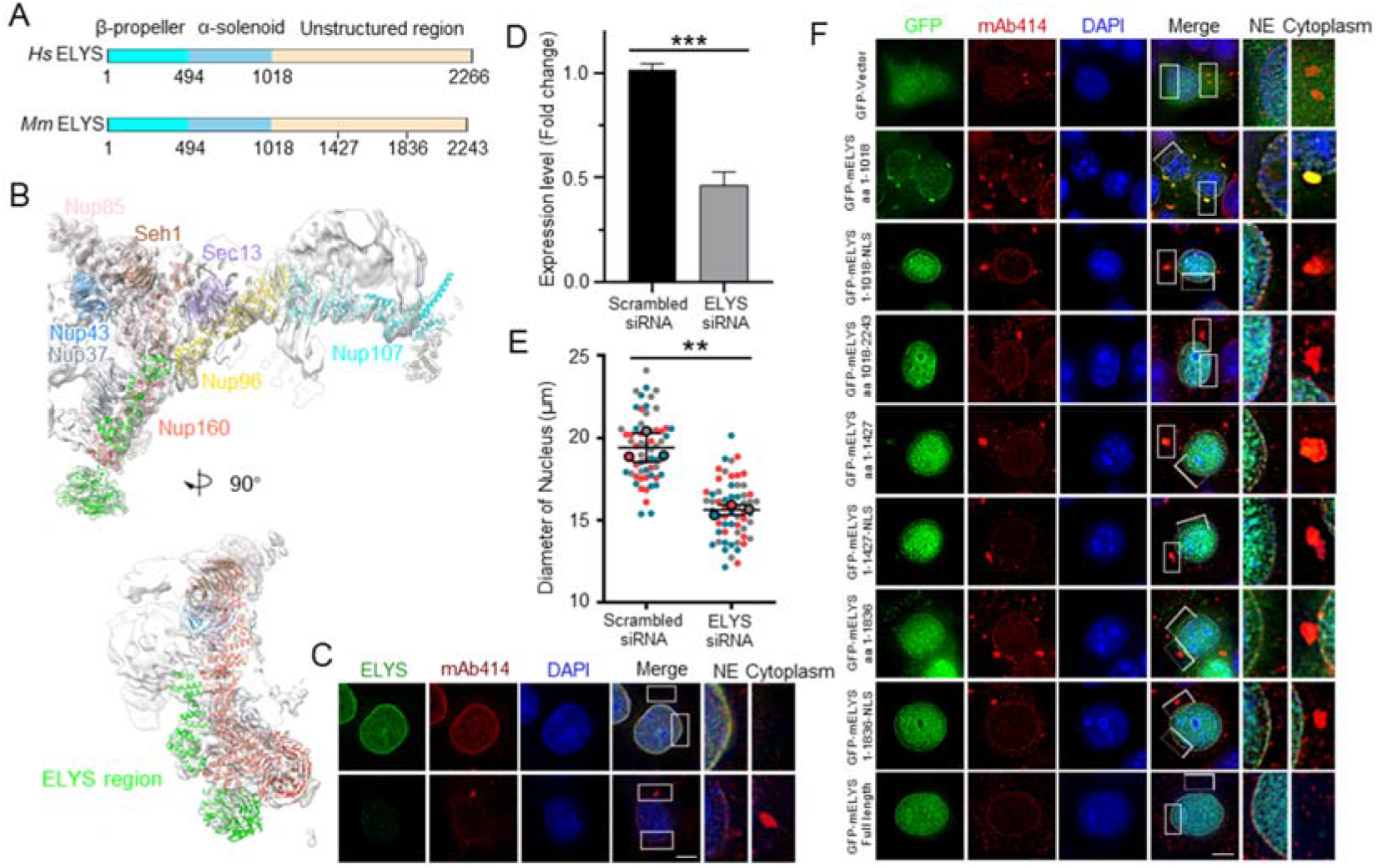
ELYS structure and function in the NPC assembly. **(A)**Organization of human (*Hs*) and mouse (*Ms*) ELYS protein. **(B)** Identification and modeling of ELYS in the inner Y complex. **(C)** RNA interference depletion of ELYS in cells. HeLa cells transfected with either scrambled or ELYS siRNA for 72 h, fixed with methanol and immunostained with specific antibodies. DNA was count-stained with DAPI. Higher magnification views of the white box areas are shown in the right panels. Scale bar, 10 μm. **(D)** Levels of ELYS mRNA detected in HeLa cells at 72 h after transfection with ELYS siRNA compared to scrambled siRNA. Error bar indicates mean ± SD. ***, p < 0.001. **(E)** Statistics of nuclear size in scrambled or ELYS siRNA knockdown cells. Three independent replicates, indicated by different colors, were carried out, and >20 cells per replicate were quantified. Error bar indicates mean ± SD. **, p < 0.01. **(F)** Immunofluorescence of cells with endogenous ELYS knockdown and expression of exogenous GFP- or GFP-tagged full-length ELYS or truncated ELYS as indicated. Scale bar, 10 μm.

The difference map between the NR and CR asymmetric units was used to identify the location of ELYS and assign its model (Fig. 3). By docking the crystal structure of the ELYS β-propeller domain (PDB code 4I0O) into the difference map directly (Fig. 5B), we clearly observed the density responsible for the structure of the α-solenoid domain (residues 689-960) of ELYS, which was modeled based on the predicted structure of *S. pombe* ELY5, a homolog of *X. laevis* ELYS (Table S1)^37^. It was reported that ELY5 binds to the NR near an interface of the Nup37-Nup120 complex (counterpart of the Nup37-Nup160 complex) (Table S1) ^38^. Although ELY5 shares only 21% identity with ELYS in the α-solenoid domain, the two proteins show patches of very conserved sequence in individual multisequence alignment (MSA). As a result, we were able to build model of the N-terminal region (residues 1-960) of ELYS by homology modeling and an iterative simulation-based refinement approach. With the model of ELYS in the Y complex, we found that its N-terminal β-propeller domain binds directly to the NE and that its adjacent α-solenoid domain forms a close contact with the α-solenoid domain of the inner Nup160 (Fig. 5B). Importantly, we only found one copy of ELYS in the asymmetric unit of the NR and observed that this ELYS interacts with the inner Nup160. Meanwhile, it’s worth noting that the local resolution in ELYS region is not high enough to assign the accurate secondary structures, so a more reliable initial model and an improved density map are required to build a more accurate pseudoatomic model for ELYS.

To further investigate the functions of ELYS, we performed knockdown experiments. We not only confirmed that ELYS is necessary for the assembly of normal NPCs on the NE, but also, interestingly, observed that knocking down ELYS resulted in significant aggregation of Nups in the cytoplasm and reduced the size of the nuclei (Fig. 5C–E). Then, we investigated the functions of different regions of ELYS by expressing GFP-tagged full-length ELYS and its truncation variants (including ELYS^1-1018^, ELYS^1-1018^-NLS (nuclear location signal), ELYS^1018-2243^, ELYS^1-1427^, ELYS^1-1427^-NLS, ELYS^1-1836^, and ELYS^1-1836^-NLS) in HeLa cells and ELYS knockdown/rescue experiments (Fig. 5 and Fig. S4). Results showed that ELYS^1-1018^ did not translocate into the nucleus properly and was largely situated on the NE/NPCs. In contrary, most ELYS^1-1018^-NLSs could both enter the nucleus and be situated on the NE/NPCs. When the ELYS truncations continued to be elongated in the CTD region, their localization signals were closer to the level of full-length ELYS, which is consistent with previous reports^39^. These results proved the importance of N-terminal structured region of ELYS for NPC assembly and its localization onto the NE.

## DISCUSSION

In this study, by utilizing integrated approaches of cryo-EM, homology modeling and cell experiment, we resolved a subnanometer resolution structure of an intact NPC scaffold from *X. laevis* oocyte NEs and successfully built a more complete model of the NR. We showed that there is only one copy of Nup205 in each asymmetric unit of the NR and that Nup205 serves as a central hub to connect three nearby Y complexes via its interactions with the inner Nup43, the outer Nup160, the outer Seh1 and the adjacent inner Nup107. Considering that both the NR and CR share a very similar architecture and conformation of the Y complex assembly, it would be interesting to discuss why there is only one copy of Nup205 in the asymmetric unit of the NR, in contrast to two copies in that of the CR. In the CR, the second Nup205 is located on the inner side of the CR and colocalizes with the Nup214 complex ^21^. We thus speculate that having this second Nup205 helps to recruit and stabilize Nup214 to form the Nup214-Nup88-Nup62 complex ^28^, which is important to facilitate the export of mRNA and therefore exists only in the CR.

Two distinct pathways for NPC assembly during the cell cycle have been found so far in metazoan cells: postmitotic NPC assembly and interphase NPC assembly. The postmitotic NPC assembly pathway initiates directly on the chromatin surface and is relatively faster than the interphase NPC assembly pathway. Postmitotic NPC assembly starts with the recruitment of ELYS to the decondensing chromatin through its DNA-binding AT-hook domain, followed by further recruitment of the Y complex and transmembrane Nup Pom121 and Nup93-205 complex to form the NPC scaffold, which is followed by further recruitment of peripheral Nups, including Nup358, Nup214 and Nup153, to complete the final maturation of the NPCs ^40–44^. Depletion of ELYS specifically interferes with postmitotic NPC assembly both in a human cell line and in a reconstituted cell-free nuclear system using *Xenopus* egg extracts ^24,41^. In comparison, Pom121 plays an early and key role in the interphase NPC assembly pathway by inserting NPCs into intact double-layered NEs ^41,45^. In this study, we found that ELYS attaches closely to the inner Nup160 at the long arm region via its α-solenoid domain and interacts with the NE via its β-propeller domain, which is consistent with its key role in recruiting Nups to chromatin to initiate NPC assembly on the NE. We also revealed a crucial role of the unstructured C-terminal region by demonstrating that it targets ELYS to the nucleus, which is also essential for initiating postmitotic NPC assembly.

In addition, we have also tried *in situ* cross linking mass spectrometry method to verify the interaction pairs in NR subunit, and revealed rich interaction pairs among NR Nups (Fig. S5A-C). For Nup205 and ELYS specifically, the results suggested their potential interactions with surrounding Nups (Fig. S5). Yet more accurate interaction patterns remained to be further investigated.

Overall, our structural study of intact NPCs from the nuclear envelope of *X. laevis* oocyte NE offers an update to the architecture and assembly of the NPCs and the construction of a subnanometer framework, advancing our understanding of nuclear transport at the molecular level.

## Materials And Methods

### Sample preparation

African clawed toad *X. laevis* maintenance, oocyte isolation, and NE preparation for cryo-EM were carried out as described previously ^8,26,46^. Briefly, ovaries were removed from narcotized mature female *X. laevis* (Nasco, USA) with a brief wash in freshly prepared amphibian Ringer’s solution (111 mM NaCl, 1.9 mM KCl, 1.1 mM CaCl2, 2.4 mM NaHCO3), and developmental stage VI oocytes were transferred to ice-cold HEPES buffer (83 mM KCl, 17 mM NaCl, 10 mM HEPES, pH 7.5) for nuclear isolation. The oocyte nuclei were isolated in HEPES buffer, applied to glow-discharged holey carbon grids (R2/1, 200 mesh, Au, Quantifoil, Germany), and the NE was spread onto the grid by fine glass needles. After careful washing in HEPES buffer, the NE on the grid was cross-linked with 0.15% glutaraldehyde in HEPES buffer on ice for 10 min. Then, for cryo-ET sample preparation, 2 μL of colloidal gold fiducial beads (10 nm diameter) was dripped onto the NE sample and allowed to rest for 1 min before plunge freezing. For cryo-EM SPA sample preparation, gold fiducial beads were not applied. The grid was blotted and vitrified by plunge freezing into liquid ethane by Vitrobot Mark IV (Thermo Fisher Scientific, USA) at 4°C and 100% humidity.

The animal experiments were performed in the Laboratory Animal Center of Peking University in accordance with the National Institutes of Health Guide for the Care and Use of Laboratory Animals and according to guidelines approved by the Institutional Animal Care and Use Committee at Peking University.

### Cryo-EM data acquisition

During the application of the NE onto the grid, the cytoplasmic ring side was kept always facing the carbon film of the grid. Prior to data collection, we first separated the grids into two groups: one group with the carbon film facing onto the C-clip of FEI AutoGrid and another group with the carbon film facing in the opposite direction. All grids were screened using a Talos Arctica 200 kV cryo-electron microscope (Thermo Fisher Scientific, USA). Then, 5971 micrographs were collected using a Titan Krios G2 300 kV cryo-electron microscope (Thermo Fisher Scientific, USA) operated in EF-TEM mode with a nominal magnification of 64,000X. The calibrated physical pixel size on the specimen was 2.24 Å. The stage tilting angles were set to 0, 30, 45 and 60 degrees. For the 0/30-degree tilting angles, the total exposure dose was set to 60 e^−^/Å^2^, with an exposure time of 21.5 seconds and 0.5 seconds per frame. For the 45-degree tilting angle, the total exposure dose was set to 80 e^−^/Å^2^, with an exposure time of 41 or 28.5 seconds for two different sessions, 0.5 seconds per frame. For the 60-degree tilting angle, the total exposure dose was set to 100 e^−^/Å^2^, with an exposure time of 36 or 35 seconds for two different sessions, 0.5 seconds per frame. The movies were acquired by a Gatan K2 Summit direct detection camera equipped with a GIF Quantum energy filter (Gatan Company, USA) with a silt width of 20 eV operated in super-resolution mode. SerialEM ^47^ with in-house scripts was used for data collection with the defocus value set between −1.0 and −4.0 μm.

### SPA image processing

The super-resolution movies were first subjected to motion correction using MotionCor2 ^48^ with a binning level of 2 in Fourier space, and dose weighting was also performed during this process. Since tilted images require accurate defocus value estimation on a per particle basis, particle picking was performed prior to CTF estimation. Particles were auto-picked using RELION-3.0 ^49^ with subsequent manual inspection. A total of 87,905 full NPC particles were selected. For 0-degree images, the defocus of each picked NPC was estimated by GCTF ^50^ using its per particle defocus estimation function. For other tilting images, the defocus of each selected NPC was estimated by goCTF ^51^ or Warp ^52^.

The overall cryo-EM SPA image processing workflow is shown in Fig. S2. The selected NPC particles were first extracted with a box size of 216 pixels and a binning level of 4, which resulted in a pixel value of 8.96 Å. With prior information on the tilting angles and the relative orientation of the grid, we were able to assign a prior tilt angle of 0/30/45/60/120/135/150/180 degrees for each particle. The refinements were performed using RELION-3.0 with the local search strategy applied. First, we ran 100 iterations of 3D classification with K = 1 using the previously reported NPC map (EMD-3103) ^9^ low-pass filtered to 60 Å as the initial reference. A symmetry of 8-fold was applied, and the tilt angle search range was restricted to 3 degrees for each iteration. Then, another round of 3D classification with K = 1 using the map reference (EMD-3103) was performed with 20 iterations and 8-fold symmetry applied. Then, we docked the previously reported model of the NR (PDB entry code 5A9Q) into the resulting map and segmented out the NR map from the whole NPC using Chimera ^53^. Based on the segmented map, we generated a local mask of the NR region. Using this mask and the output star file from the previous round of 3D classification, we performed auto-refinement to obtain a 27 Å resolution map of the NR with the 8-fold symmetry applied.

With the refined shifts and orientations of the NPC particles, we re-extracted particles with a box size of 400 pixels and binned pixel size of 4.48 Å. Using a similar strategy as that above, we reconstructed the cryo-EM map of the NR at a resolution of 22 Å. With this better resolution, we achieved a more accurate determination of each NR asymmetric unit. We used Chimera to measure the relative coordinates of one asymmetric unit of the NR and generated a symmetry expanded particle star file with updated defocus corresponding to each NR asymmetric unit by using a modified version of a block-based reconstruction script (Script S1) ^27^. Then, we re-extracted particles with a box size of 200 pixels and a binning level of 2. Another round of auto-refinement was performed using the generated star file and the NR asymmetric unit mask, yielding the cryo-EM map of the NR asymmetric unit at a resolution of 13 Å.

With the refined shifts and orientations of each asymmetric unit, we re-extracted particles with a box size of 320 pixels, binning level of 1 and pixel size of 2.24 Å. We first performed a reconstruction to ensure that all the predetermined parameters were correct and to generate a better mask containing one asymmetric unit of the NR. Then, another round of auto-refinement yielded a map at a resolution of 10 Å. Next, we ran a 3D classification job with a T value of 10.

Particles corresponding to the best class were selected, and refinement of these particles reached a resolution of 7.8 Å. A stable core region was identified by investigating the local resolution distribution of this 7.8 Å map, and the mask covering this region only was created. The 3D classification and refinement of the stable core region were performed using the polished data, the 7.8 Å map as the reference, and the corresponding mask, which resulted in a final resolution of 6.8 Å after postprocessing. A similar strategy was applied to the UDR of the NR and resulted in a resolution of 10.6 Å. The similar image processing approach was applied to the asymmetric unit of IR and CR, which yielded a resolution of 9.8 and 7.6 Å after postprocessing in RELION-3.0 (Table S2). All figures in this work were generated by Chimera ^53^ or ChimeraX ^54^.

### Calculation of the difference map

The difference maps between asymmetric units of CR and NR were calculated using EMAN2 ^55^. First, the cryo-EM map of CR asymmetric unit was fitted into that of NR and then rescaled onto the same coordinate system using the command line tool (*vop resample #1 ongrid #2*) in Chimera ^53^. Then the structural amplitudes of both maps were scaled using EMAN2. Finally, the difference maps were computed using EMAN2, simply by multiplying −1 to one map and then add it onto another.

### Cryo-ET data acquisition and processing

For cryo-ET data acquisition, a total of 334 tilt series were collected on a Titan Krios G2 300 kV cryo-electron microscope (Thermo Fisher Scientific, USA) operated in EF-TEM mode with a nominal magnification of 42000x, which resulted in a calibrated physical pixel size of 3.4 Å at the specimen level. Tilt series between −60 ° and +60° with a 3° angular step using a dose-symmetric scheme were acquired ^56^. A total dose of 143.5 e^−^/Å^2^ per tilt series was distributed evenly among 41 tilts, and the defocus value was set between −1.5 and −3.0 μm.

Then, we used Warp ^52^ to perform the preprocessing, including motion correction and picking and masking of fiducial markers. After preprocessing, all tilt series stacks were generated using automatic procedures in Warp. Alignment of tilt series and transformation of alignment file formats were performed using a wrapped package ^57^ of automatic tilt series alignment functions in the Dynamo ^58^ and IMOD ^59^ packages. The alignment files were transferred back to Warp to perform per-tilt CTF estimation. Tomograms were reconstructed in Warp at a binning level of 8 with a pixel size of 27.2 Å. Particle picking was performed using template matching functions in Dynamo with the map reference (EMD-3103) low-pass filtered to 80 Å. After the template matching process, manual inspections using Dynamo were performed for all the tomograms to further validate particle picking accuracy. In total, 1360 particles were extracted from Warp reconstructed tomograms using Dynamo with a box size of 72 pixels, a reference was generated by direct averaging of extracted particles, and alignment was performed for 4 iterations with 8-fold symmetry applied in Dynamo. Then, the aligned coordinates and orientations were transferred back to Warp for re-extraction of 4 binned particles with a box size of 144 pixels and pixel size of 13.6 Å. Further refinement was performed using RELION-3.0 with 8-fold symmetry applied (Table S3).

### Homology modeling

Considering the sequences of *X. laevis* Nup160 and Nup96 are not available from the public databases, we utilized the sequences of their homologues from *Xenopus tropicalis* (*X. tropicalis*) to build the models.

The structure of the NR Y complexes was resolved by iteratively combining homology modeling and simulation-aided structure refinement. The NR Y complex was divided into two regions according to the quality of the collected EM densities: the core region with the highest resolution and the peripheral region with relatively lower local resolutions. The core region was mainly composed of Nup85, Nup96, Nup160 CTD (aa. 874-1432), Seh1, Nup43, Sec13, and Nup37. In addition to homology modeling using previously resolved crystallographic structures as templates, there were segments lacking either a structural template or a considerable topological similarity to the template. We therefore used the GALAXY template-based modeling program ^60^ to model the missing segments, including aa. 473-646 of Nup85, aa. 736-862 of Nup96, and aa. 873-1432 of Nup160. The relative orientation between the beta propellers (Seh1, Nup43, Sec13 and Nup37) and structural arms formed by Nup85, Nup96 and Nup160 was determined by structural comparison to previously resolved oligomeric structures as follows: PDB-3F3F for the Seh1-Nup85 complex, PDB-3BG1 for the Sec13-Nup96 complex, and PDB-4GQ2 for the Nup37-Nup160 complex. Although an oligomeric structure of the Nup43-Nup85 complex is lacking, we oriented Nup43 in accordance with the Nup37-Nup160 complex based on its distal topological features compared to other beta propellers found in the NR Y complex. We then performed stepwise MDFF ^61–63^ simulations of the core region in which the collected EM densities were converted to biasing potentials added to standard molecular dynamics (MD) simulations. We gradually reduced the scaling factor, which dictates the strength of the bias from 0.5 to 0.1 during the simulation to allow more aggressive relaxation of the protein-protein interfaces and to avoid overfitting.

The peripheral regions referred to proteins branching from the central arms or at the interface of two Y complexes, including the Nup160 NTD (aa. 45-436), Nup107, Nup133, Nup205, and ELYS. As the resolution of the EM densities for the peripheral regions was lower than that of the core region, we employed an iterative approach to achieve atomistic structure refinement of the protein complexes. We first performed homology modeling for individual Nup proteins or protein segments, including the Nup107 NTD (aa. 142-590), Nup107 CTD (aa. 658-915), Nup133 beta propeller (aa. 9-407), Nup133 alpha solenoid NTD (aa. 518-888), Nup133 alpha solenoid CTD (aa. 905-1139), Nup205 NTD (aa. 39-498), Nup205 central segment (aa. 499-951), Nup205 CTD1 (aa. 952-1409), Nup205 CTD2 (aa. 1410-1691), ELYS beta propeller (aa. 3-491), and ELYS alpha solenoid (aa. 689-960). We therefore used the GALAXY template-based modeling program ^60^ to model the missing segments (modeled regions), including aa. 591-657 of Nup107, aa. 889-904 of Nup133, and aa. 1-38 of Nup205. The following structural comparisons were used to determine the relative orientation in the protein complexes: PDB-3IKO for Nup107-Nup96 and PDB-3I4R for Nup133-Nup107. The protein complexes were then subjected to MDFF simulations, including components from the core region if they constituted a protein-protein interface, to flexibly relax the modeled regions. To determine a convergence in the modeled structure, we measured the backbone root-mean-square deviation (RMSD) from the modeled structure before refinement and set the threshold to 1 Å. Iteratively, all the side chains in the modeled regions and protein-protein interfaces were remodeled using MODELLER ^64^ and subjected to a new round of structural refinement through MDFF simulations.

Homology modeling was performed using SWISS-MODEL ^65^, the GALAXY template-based modeling program ^60^, and MODELLER ^64^ with multisequence alignment (MSA) obtained using the HH suite ^66^ on the latest Uniclust30 database at the time (UniRef30_2020_06) ^67^. Consensus among the methods was obtained. The secondary structure predictions were performed using PSIPRED ^68^.

### Structure determination of ELYS

Although ELY5 shares only 21% identity with the ELYS domain of ELYS, the two proteins show very common conserved sequence patches from individual MSAs. We then performed homology modeling of the β-propeller and the ELYS domain. Residues 492-688 were omitted because MSA revealed no query with sequence coverage larger than 30%. The orientation of ELYS with respect to Nup160 was optimized with local redocking using HADDOCK ^69^ before it was flexibly fitted to the differential maps. Again, the backbone coordinates of the modeled structure were extensively refined by the iterative MDFF protocol described above.

### MDFF simulations

Before MDFF simulation, the protein complex was solvated in explicit TIP3P water molecules ^70^ with sodium and chloride ions at a final concentration of 0.15, and steepest descent energy minimization was performed on the initial structure for at least 10,000 steps before the refinement run. A timestep of 1 fs was used throughout the simulation. The refinement runs were performed for 100 to 500 ps, which corresponds to 100,000 to 500,000 simulation steps depending on the convergence of the cross-correlation coefficient (CCC) profile. The scaling factor was decreased from 0.5 by 0.1 decrements every 20,000 simulation steps. All simulations were performed using CHARMM36m ^71^ forcefields. Langevin dynamics were adopted at a temperature of 310 K. The Nose-Hoover Langevin-piston method ^72^ was used in the constant pressure simulations, with a targeted pressure at 1 atm. Electrostatic calculations were treated with particle mesh Ewald (PME) ^73^. A cutoff of 12 Å was chosen for short-range van der Waals interactions. NAMD ^74^ was used as the MD engine throughout all simulations.

### Crosslinking Mass Spectrometry

The nuclei were isolated from HeLa cells with nuclear extraction reagents (Thermo, NE-PER™) and suspended in 1 X PBS buffer (pH 7.4). The suspension of intact nuclei was crosslinked by adding 5 mM BSP (Fig. S4), having the maximum Cα-Cα distance restraints of 28 Å, every 20 min for 3 times at room temperature and quenched with 50 mM ammonium bicarbonate. The sample was adjusted to 1% SDS and sonicated until nuclei were lysed. Then the lysate was diluted to 0.2% SDS and subjected to the click chemistry reaction with diazo biotin-azide enrichment reagent. Afterwards, the chemically cross-linked sample was precipitated using cold acetone and performed the subsequent proteolytic digestion.

To reduce the abundance suppression of histone complex on nuclear ring protein complex (NRPC), the sample was solubilized in 8M urea, then transferred into the 0.1 μm ultrafiltration device to retain the high-molecular-weight NRPC and to allow through low-molecular–weight histone complex. Then a 300 K ultrafiltration device was adopted to retrieve the potential missing NRPC from filtrate. After that, the sample retained on the filter was respectively reduced by 10 mM tris-(2-carboxyethyl)-phosphine for 1h at room temperature, and alkylated by 20 mM iodoacetamide for 20 min in the dark. Then the cross-linked complex was digested with trypsin at ratio of 1:20 (enzyme/protein, w/w) to generate the cross-linked peptides. Finally, the cross-linked peptides were enriched with streptavidin beads for 2 h at room temperature, and released from the beads by Na2S2O4 buffer, followed by LC-MS/MS analysis.

To further improve the identification coverage of NRPC, the cross-linker of DMTMM was used, combining with the amino-reactive BSP for the crosslinking of NEs isolated from HeLa cells with commercial kit (Invent, NE-013™), respectively. As a supplementary, the carboxyl group of Glu and Asp can be activated by DMTMM to form an active O-acylisourea intermediate that would react with a spatially adjacent amino group of Lys to yield a stable imide linkage under physiological conditions, thus increasing the identification coverage of crosslinking sites.

To reduce the complexity of the cross-linked peptides, high pH RPLC was used to separate the peptides into 6–15 fractions with a C18 column (5 μm, 100 Å, 150 mm × 2.1 mm i.d.). Each peptide fraction was dissolved in the sample loading buffer (0.1% FA) and analyzed by LC-MS/MS using an Easy-nano LC 1200 system connected online to an Orbitrap Fusion Lumos mass spectrometer (Thermo). Samples were automatically loaded onto a C18 RP trap column (150 μm i.d. × 3 cm) and separated by a C18 capillary column (150 μm i.d. × 15 cm), packed in-house with ReproSil-Pur C18-AQ particles (1.9 μm, 120 Å) using a stepwise 100 min gradient between 6 and 40% (v/v) ACN in 0.1% (v/v) FA. The mass spectrometer was operated in positive ion mode. Full MS scans were acquired in the orbitrap analyzer over the m/z 350-1500 range with a resolution of 60000 and the AGC target was 4e5. Peptides (charge states from 3+ to 7+) were selected for subsequent MS/MS scans with a resolution of 15000. The maximum allowed ion accumulation times were 50 ms for MS scans and 30 ms for MS/MS scans. The raw data were searched by pLink2 software using a FASTA database containing 22 gene-coding nuclear ring protein sequences. The data were automatically filtered using the mass error of 20 ppm for precursor ions and fragment ions, respectively. Other search parameters included cysteine carbamidomethyl as a fixed modification and the oxidation of methionine and the acetylation of protein N termini as a variable modification. A maximum of three trypsin missed-cleavage sites was allowed. The minimum peptide length was specified to five amino acids, and the FDR ≤ 1% at PSM level was set to control the data threshold separately.

Besides, PRM analysis was used to verify the identified cross-linked peptides of NRPC. Distribution of Cα–Cα distances of the cross-linked sites identified were validated by the PDB structure of nuclear pore complex (PDB code: 3TJ3, 4LIR, 5A9Q, 5IJN, 5IJO, 5TO5). The maximum distance restraint imposed by BSP is 28 Å.

### In-lens field emission scanning electron microscopy (In-Lens FESEM)

NE from *X. laevis* oocytes was spread onto silicon chips as described previously ^26^ and then fixed with 2% glutaraldehyde in 0.1 M sodium cacodylate buffer (pH 7.4) at room temperature for 30 min. The sample was rinsed and postfixed with 1.0% OsO4 in 0.1 M sodium cacodylate buffer at room temperature for 30 min. After dehydration in a graded ethanol series, the sample was transferred to a CO2 critical point dryer. The sample was then coated with 5 nm gold in a Hitachi E-1045 ION sputter and viewed in a Hitachi In-lens field emission scanning electron microscope S-4800 (Hitachi, Japan) at an accelerating voltage of 4 kV.

### Transmission electron microscopy ultrathin sections

For TEM ultrathin sections, the NE of *X. laevis* oocytes was spread onto 35 mm cell culture dishes and fixed with 2.5% glutaraldehyde in 0.1 M sodium cacodylate buffer (pH 7.4) at room temperature for 30 min. The sample was rinsed and postfixed with 1.0% OsO4 in 0.1 M sodium cacodylate buffer at room temperature for 30 min. Then, the sample was rinsed and stained in 1% aqueous uranyl acetate for 20 min. After dehydration of a graded series of ethanol, the sample was embedded in Epon-812 resin and sectioned with a diamond knife and a Leica Ultracut R cutter. After staining with aqueous uranyl acetate and lead citrate, the sections were observed under an FEI Tecnai G2 20 Twin TEM (Thermo Fisher Scientific, USA), and images were captured with an Eagle 4K CCD camera (Thermo Fisher Scientific, USA) ^75^.

### Immunofluorescence microscopy (IFM)

HeLa cells were grown onto glass coverslips in Dulbecco’s modified Eagle’s medium (DMEM) with 10% fetal calf serum at 37°C in a 5% CO2 atmosphere and transfected with appropriate plasmids. After 24 h, cells were fixed in methanol for 5 min on ice, stained with ELYS antibody (Novus Biologicals, NBP1-87952) or mAb414 antibody (Covance, MMS-120P) and 4’6’-diamidino-2-phenylindole (DAPI) (Sigma-Aldrich, D9542), and then observed with a Delta Vision Elite fluorescence microscope (GE, USA).

### RNA interference and rescue

To knock down ELYS *in vivo*, chemically synthesized siRNAs were used. The siRNA sequences were as follows: ELYS siRNA, 5’-GGAACUGUGUUGACAAGAUTT-3’; scrambled siRNA negative control, 5’-UUCUCCGAACGUGUCACGUTT-3’. HeLa cells were transfected using 100 pmol of siRNA with 5 μL of Lipofectamine 2000 (Invitrogen, 11668019) for 72 h ^76^. The cells were then collected for IFM and counted for statistics. For rescue experiments, since we failed to acquire the full-length DNA sequence of the human ELYS gene, and since the human and mouse ELYS proteins share high sequence homology (76.7% similarity), the mouse ELYS gene was used in plasmid construction, including GFP-C1 vector as a negative control, full-length mouse ELYS as a positive control and multiple ELYS truncations (residues 1-1018, 1-1018-NLS, 1018-2243, 1-1427, 1-1427-NLS, 1-1836, 1-1836-NLS). The NLS sequence we used was GTCACCAAAAAGCGCAAACTGGAGTCCACT. HeLa cells were first transfected with siRNA for 24 h and then transfected with different plasmids by Lipofectamine 2000 (Invitrogen, 11668019) again. After 48 h, the cells were collected for IFM.

### RNA isolation, cDNA synthesis and quantitative PCR

Since our ELYS antibody could not effectively detect ELYS protein by western blot, we used quantitative PCR to detect the interference efficiency of ELYS siRNA. HeLa cells were transfected with scrambled and ELYS siRNAs for 72 h. Then, total RNA was isolated from the cells using TRIzol reagent (Invitrogen, 15596026) according to the recommendations of the manual. One microgram of total RNA was reverse transcribed by the PrimeScript RT Reagent Kit with gDNA Eraser (Takara, RR047A). Quantitative PCR was performed in technical duplicates with FastStart Essential DNA Green Master Mix (Roche, 06402712001) and a LightCycler 96 instrument (Roche, 05815916001). Quantification results were analyzed by LightCycler Software Version 1.1.0. Gene expression levels were normalized to the housekeeping gene β-actin. Specific primers were as follows: β-actin forward/reverse: TCGTGCGTGACATTAAGGAG / GTCAGGCAGCTCGTAGCTCT; ELYS forward/reverse: GCAGCAGCAGGACTCGGTCT / TCCTTGGAACTTCTGACGCTGGA, as reported ^77^.

## Data Availability

The Electron Microscopy Database (EMD) accession codes of the NR asymmetric unit, NR stable core region, map containing UDR, CR asymmetric unit and IR asymmetric unit are EMD-31939, EMD-31940, EMD-31941, EMD-31942, EMD-31943, respectively. The Protein Data Bank (PDB) accession code of the model of the NR asymmetric unit is PDB 7VE1. The raw MS data files of crosslinking proteomics have been deposited to the integrated proteome resources (iProX) with project ID IPX0003500000.

## Acknowledgements

We thank all other members of the Fei Sun and Chuanmao Zhang laboratories for their help and critical comments and all other members of the Lihua Zhang laboratory for their help with CX-MS. We would also like to thank the Center for Biological Imaging (CBI), Institute of Biophysics, Chinese Academy of Science for cryo-EM work and Boling Zhu, Xujing Li and Gang Ji for their help with cryo-EM data collection; the Facilities Cores at National Center for Protein Sciences and the cryo-EM and TEM platforms at the College of Life Sciences of Peking University for cryo-electron microscopy and TEM; and Ning Gao, Zhenxi Guo, Guopeng Wang, Yingchun Hu, Xia Pei and Bo Shao for their help with cryo-EM and TEM experiments. We thank Mengqiu Dong and Yong Cao (National Institute of Biological Sciences, Beijing, China) for their help at the early stage of the project for cross-linking mass spectrometry experiments. This work was equally supported by grants from Ministry of Science and Technology of China (2017YFA0504700 to FS and 2016YFA0500201 to CMZ), the Strategic Priority Research Program of the Chinese Academy of Sciences (XDB 37040102 to FS), and National Natural Science Foundation of China (31830020 to FS, 31520103906 to CMZ, and 2018YFA050703 to QZ). This work was also supported by grants from the National Science Fund for Distinguished Young Scholars (31925026 to FS and 21725506 to LZ), National Natural Science Foundation of China (31430051 to CMZ) and National Key Research and Development Program of China (2016YFA0100501 to CMZ and 2018YFA0901102 to YZ). This work was also supported by the Research Grant Council of Hong Kong (11306517, 11305919, and 11308620 to JF) and the National Science Foundation of China/RGC Joint Research Scheme (N_CityU104/19 to JF).

## AUTHOR CONTRIBUTIONS

C. Z. and F. S. conceived the project and designed the experiments. H. R., L. T. and X. H. performed cryo-EM experiments. L. T., Y. Z., J. X. and F. S. performed cryo-EM data processing. H. R., L. T., M. J., G. Z. and X. W. participated in the preparation and screening of cryo-EM samples. C. C., J. F., X. M., X. Z., and C. H. performed the homology modeling and simulation-based refinement. LH. Z., Q. Z., LL. Z., Y. A. and G. Y. performed CX-MS experiments. H.R. performed the ELYS mutagenesis study. H. R., L. T., Y. Z., C. C. and Q. Z. analyzed the data and wrote the manuscript, which was substantially revised by F. S. and C. Z.

## References

1 Clarke, P. R. & Zhang, C. Spatial and temporal coordination of mitosis by Ran GTPase. Nat Rev Mol Cell Biol 9, 464–477, doi:10.1038/nrm2410 (2008).

2 Lin, D. H. & Hoelz, A. The Structure of the Nuclear Pore Complex (An Update). Annu Rev Biochem 88, 725–783, doi:10.1146/annurev-biochem-062917-011901 (2019).

3 Kim, S. J. et al. Integrative structure and functional anatomy of a nuclear pore complex. Nature 555, 475–482, doi:10.1038/nature26003 (2018).

4 Grossman, E., Medalia, O. & Zwerger, M. Functional architecture of the nuclear pore complex. Annu Rev Biophys 41, 557–584, doi:10.1146/annurev-biophys-050511-102328 (2012).

5 Lin, D. H. et al. Architecture of the symmetric core of the nuclear pore. Science 352, aaf1015, doi:10.1126/science.aaf1015 (2016).

6 Callan, H. G. & Tomlin, S. G. Experimental studies on amphibian oocyte nuclei. I. Investigation of the structure of the nuclear membrane by means of the electron microscope. Proc R Soc Lond B Biol Sci 137, 367–378, doi:10.1098/rspb.1950.0047 (1950).

7 Bui, K. H. et al. Integrated structural analysis of the human nuclear pore complex scaffold. Cell 155, 1233–1243, doi:10.1016/j.cell.2013.10.055 (2013).

8 Eibauer, M. et al. Structure and gating of the nuclear pore complex. Nat Commun 6, 7532, doi:10.1038/ncomms8532 (2015).

9 von Appen, A. et al. In situ structural analysis of the human nuclear pore complex. Nature 526, 140–143, doi:10.1038/nature15381 (2015).

10 Knockenhauer, K. E. & Schwartz, T. U. The Nuclear Pore Complex as a Flexible and Dynamic Gate. Cell 164, 1162–1171, doi:10.1016/j.cell.2016.01.034 (2016).

11 Hsia, K. C., Stavropoulos, P., Blobel, G. & Hoelz, A. Architecture of a coat for the nuclear pore membrane. Cell 131, 1313–1326, doi:10.1016/j.cell.2007.11.038 (2007).

12 Brohawn, S. G., Leksa, N. C., Spear, E. D., Rajashankar, K. R. & Schwartz, T. U. Structural evidence for common ancestry of the nuclear pore complex and vesicle coats. Science 322, 1369–1373, doi:10.1126/science.1165886 (2008).

13 Debler, E. W. et al. A fence-like coat for the nuclear pore membrane. Mol Cell 32, 815–826, doi:10.1016/j.molcel.2008.12.001 (2008).

14 Seo, H. S. et al. Structural and functional analysis of Nup120 suggests ring formation of the Nup84 complex. P Natl Acad Sci USA 106, 14281–14286, doi:10.1073/pnas.0907453106 (2009).

15 Berke, I. C., Boehmer, T., Blobel, G. & Schwartz, T. U. Structural and functional analysis of Nup133 domains reveals modular building blocks of the nuclear pore complex. J Cell Biol 167, 591–597, doi:10.1083/jcb.200408109 (2004).

16 Boehmer, T., Jeudy, S., Berke, I. C. & Schwartz, T. U. Structural and functional studies of Nup107/Nup133 interaction and its implications for the architecture of the nuclear pore complex. Mol Cell 30, 721–731, doi:10.1016/j.molcel.2008.04.022 (2008).

17 Kampmann, M. & Blobel, G. Three-dimensional structure and flexibility of a membrane-coating module of the nuclear pore complex. Nat Struct Mol Biol 16, 782–788, doi:10.1038/nsmb.1618 (2009).

18 Stuwe, T. et al. Architecture of the fungal nuclear pore inner ring complex. Science 350, 56–64, doi:10.1126/science.aac9176 (2015).

19 Beck, M. et al. Nuclear pore complex structure and dynamics revealed by cryoelectron tomography. Science 306, 1387–1390, doi:10.1126/science.1104808 (2004).

20 Zhang, Y. et al. Molecular architecture of the luminal ring of the Xenopus laevis nuclear pore complex. Cell Res, doi:10.1038/s41422-020-0320-y (2020).

21 Huang, G. et al. Structure of the cytoplasmic ring of the Xenopus laevis nuclear pore complex by cryo-electron microscopy single particle analysis. Cell Res, doi:10.1038/s41422-020-0319-4 (2020).

22 Allegretti, M. et al. In-cell architecture of the nuclear pore and snapshots of its turnover. Nature 586, 796–800, doi:10.1038/s41586-020-2670-5 (2020).

23 Kosinski, J. et al. Molecular architecture of the inner ring scaffold of the human nuclear pore complex. Science 352, 363–365, doi:10.1126/science.aaf0643 (2016).

24 Franz, C. et al. MEL-28/ELYS is required for the recruitment of nucleoporins to chromatin and postmitotic nuclear pore complex assembly. Embo Rep 8, 165–172, doi:10.1038/sj.embor.7400889 (2007).

25 Aebi, U., Cohn, J., Buhle, L. & Gerace, L. The Nuclear Lamina Is a Meshwork of Intermediate-Type Filaments. Nature 323, 560–564, doi:10.1038/323560a0 (1986).

26 Allen, T. D. et al. A protocol for isolating Xenopus oocyte nuclear envelope for visualization and characterization by scanning electron microscopy (SEM) or transmission electron microscopy (TEM). Nat Protoc 2, 1166–1172, doi:10.1038/nprot.2007.137 (2007).

27 Zhu, D. et al. Pushing the resolution limit by correcting the Ewald sphere effect in single-particle Cryo-EM reconstructions. Nat Commun 9, 1552, doi:10.1038/s41467-018-04051-9 (2018).

28 Huang, G. et al. Structure of the cytoplasmic ring of the Xenopus laevis nuclear pore complex by cryo-electron microscopy single particle analysis. Cell Res 30, 520–531, doi:10.1038/s41422-020-0319-4 (2020).

29 Mosalaganti, S. et al. In situ architecture of the algal nuclear pore complex. Nat Commun 9, 2361, doi:10.1038/s41467-018-04739-y (2018).

30 Liebschner, D. et al. Macromolecular structure determination using X-rays, neutrons and electrons: recent developments in Phenix. Acta Crystallogr D 75, 861–877, doi:10.1107/S2059798319011471 (2019).

31 Drin, G. et al. A general amphipathic alpha-helical motif for sensing membrane curvature. Nat Struct Mol Biol 14, 138–146, doi:10.1038/nsmb1194 (2007).

32 Hampoelz, B., Andres-Pons, A., Kastritis, P. & Beck, M. Structure and Assembly of the Nuclear Pore Complex. Annu Rev Biophys 48, 515–536, doi:10.1146/annurev-biophys-052118-115308 (2019).

33 Rasala, B. A., Orjalo, A. V., Shen, Z. X., Briggs, S. & Forbes, D. J. ELYS is a dual nucleoporin/kinetochore protein required for nuclear pore assembly and proper cell division. P Natl Acad Sci USA 103, 17801–17806, doi:10.1073/pnas.0608484103 (2006).

34 Gillespie, P. J., Khoudoli, G. A., Stewart, G., Swedlow, J. R. & Blow, J. J. ELYS/MEL-28 chromatin association coordinates nuclear pore complex assembly and replication licensing. Curr Biol 17, 1657–1662, doi:10.1016/j.cub.2007.08.041 (2007).

35 Ren, H. et al. Postmitotic annulate lamellae assembly contributes to nuclear envelope reconstitution in daughter cells. J Biol Chem 294, 10383–10391, doi:10.1074/jbc.AC119.008171 (2019).

36 Bilokapic, S. & Schwartz, T. U. Structural and functional studies of the 252 kDa nucleoporin ELYS reveal distinct roles for its three tethered domains. Structure 21, 572–580, doi:10.1016/j.str.2013.02.006 (2013).

37 Zimmerli, C. E. et al. Nuclear pores constrict upon energy depletion. bioRxiv, 2020.2007.2030.228585, doi:10.1101/2020.07.30.228585 (2020).

38 Bilokapic, S. & Schwartz, T. U. Molecular basis for Nup37 and ELY5/ELYS recruitment to the nuclear pore complex. Proc Natl Acad Sci U S A 109, 15241–15246, doi:10.1073/pnas.1205151109 (2012).

39 Gomez-Saldivar, G. et al. Identification of Conserved MEL-28/ELYS Domains with Essential Roles in Nuclear Assembly and Chromosome Segregation. PLoS Genet 12, e1006131, doi:10.1371/journal.pgen.1006131 (2016).

40 Imamoto, N. & Funakoshi, T. Nuclear pore dynamics during the cell cycle. Curr Opin Cell Biol 24, 453–459, doi:10.1016/j.ceb.2012.06.004 (2012).

41 Doucet, C. M., Talamas, J. A. & Hetzer, M. W. Cell Cycle-Dependent Differences in Nuclear Pore Complex Assembly in Metazoa. Cell 141, 1030–1041, doi:10.1016/j.cell.2010.04.036 (2010).

42 Hampoelz, B., Andres-Pons, A., Kastritis, P. & Beck, M. Structure and Assembly of the Nuclear Pore Complex. Annual Review of Biophysics, Vol 48 48, 515–536, doi:10.1146/annurev-biophys-052118-115308 (2019).

43 Zhang, C. & Clarke, P. R. Chromatin-independent nuclear envelope assembly induced by Ran GTPase in Xenopus egg extracts. Science 288, 1429–1432, doi:10.1126/science.288.5470.1429 (2000).

44 Zhang, C. & Clarke, P. R. Roles of Ran-GTP and Ran-GDP in precursor vesicle recruitment and fusion during nuclear envelope assembly in a human cell-free system. Curr Biol 11, 208–212, doi:10.1016/s0960-9822(01)00053-7 (2001).

45 Lu, Q. et al. Chromatin-bound NLS proteins recruit membrane vesicles and nucleoporins for nuclear envelope assembly via importin-alpha/beta. Cell Res 22, 1562–1575, doi:10.1038/cr.2012.113 (2012).

46 Akey, C. W. & Radermacher, M. Architecture of the Xenopus nuclear pore complex revealed by three-dimensional cryo-electron microscopy. J Cell Biol 122, 1–19, doi:10.1083/jcb.122.1.1 (1993).

47 Mastronarde, D. N. Automated electron microscope tomography using robust prediction of specimen movements. J Struct Biol 152, 36–51, doi:10.1016/j.jsb.2005.07.007 (2005).

48 Zheng, S. Q. et al. MotionCor2: anisotropic correction of beam-induced motion for improved cryo-electron microscopy. Nat Methods 14, 331–332, doi:10.1038/nmeth.4193 (2017).

49 Zivanov, J. et al. New tools for automated high-resolution cryo-EM structure determination in RELION-3. Elife 7, doi:10.7554/eLife.42166 (2018).

50 Zhang, K. Gctf: Real-time CTF determination and correction. J Struct Biol 193, 1–12, doi:10.1016/j.jsb.2015.11.003 (2016).

51 Su, M. goCTF: Geometrically optimized CTF determination for single-particle cryo-EM. J Struct Biol 205, 22–29, doi:10.1016/j.jsb.2018.11.012 (2019).

52 Tegunov, D. & Cramer, P. Real-time cryo-electron microscopy data preprocessing with Warp. Nat Methods 16, 1146–1152, doi:10.1038/s41592-019-0580-y (2019).

53 Pettersen, E. F. et al. UCSF chimera - A visualization system for exploratory research and analysis. J Comput Chem 25, 1605–1612, doi:10.1002/jcc.20084 (2004).

54 Pettersen, E. F. et al. UCSF ChimeraX: Structure visualization for researchers, educators, and developers. Protein Sci 30, 70–82, doi:10.1002/pro.3943 (2021).

55 Tang, G. et al. EMAN2: An extensible image processing suite for electron microscopy. J Struct Biol 157, 38–46, doi:10.1016/j.jsb.2006.05.009 (2007).

56 Hagen, W. J. H., Wan, W. & Briggs, J. A. G. Implementation of a cryo-electron tomography tilt-scheme optimized for high resolution subtomogram averaging. J Struct Biol 197, 191–198, doi:10.1016/j.jsb.2016.06.007 (2017).

57 Burt, A., Gaifas, L., Dendooven, T. & Gutsche, I. Tools enabling flexible approaches to high-resolution subtomogram averaging. bioRxiv doi:https://doi.org/10.1101/2021.01.31.428990 (2021).

58 Castano-Diez, D., Kudryashev, M., Arheit, M. & Stahlberg, H. Dynamo: A flexible, user-friendly development tool for subtomogram averaging of cryo-EM data in high-performance computing environments. J Struct Biol 178, 139–151, doi:10.1016/j.jsb.2011.12.017 (2012).

59 Kremer, J. R., Mastronarde, D. N. & McIntosh, J. R. Computer visualization of three-dimensional image data using IMOD. J Struct Biol 116, 71–76, doi:10.1006/jsbi.1996.0013 (1996).

60 Ko, J., Park, H., Heo, L. & Seok, C. GalaxyWEB server for protein structure prediction and refinement. Nucleic Acids Res 40, W294–297, doi:10.1093/nar/gks493 (2012).

61 Chan, K. Y. et al. Symmetry-restrained flexible fitting for symmetric EM maps. Structure 19, 1211–1218, doi:10.1016/j.str.2011.07.017 (2011).

62 Trabuco, L. G., Villa, E., Mitra, K., Frank, J. & Schulten, K. Flexible fitting of atomic structures into electron microscopy maps using molecular dynamics. Structure 16, 673–683, doi:10.1016/j.str.2008.03.005 (2008).

63 Trabuco, L. G., Villa, E., Schreiner, E., Harrison, C. B. & Schulten, K. Molecular dynamics flexible fitting: a practical guide to combine cryo-electron microscopy and X-ray crystallography. Methods 49, 174–180, doi:10.1016/j.ymeth.2009.04.005 (2009).

64 Eswar, N., Eramian, D., Webb, B., Shen, M. Y. & Sali, A. Protein structure modeling with MODELLER. Methods Mol Biol 426, 145–159, doi:10.1007/978-1-60327-058-8_8 (2008).

65 Biasini, M. et al. SWISS-MODEL: modelling protein tertiary and quaternary structure using evolutionary information. Nucleic Acids Res 42, W252–258, doi:10.1093/nar/gku340 (2014).

66 Steinegger, M. et al. HH-suite3 for fast remote homology detection and deep protein annotation. BMC bioinformatics 20, 473, doi:10.1186/s12859-019-3019-7 (2019).

67 Mirdita, M. et al. Uniclust databases of clustered and deeply annotated protein sequences and alignments. Nucleic Acids Res 45, D170–d176, doi:10.1093/nar/gkw1081 (2017).

68 Buchan, D. W. A. & Jones, D. T. The PSIPRED Protein Analysis Workbench: 20 years on. Nucleic acids research 47, W402–w407, doi:10.1093/nar/gkz297 (2019).

69 van Zundert, G. C. P. et al. The HADDOCK2.2 Web Server: User-Friendly Integrative Modeling of Biomolecular Complexes. J Mol Biol 428, 720–725, doi:10.1016/j.jmb.2015.09.014 (2016).

70 Jorgensen, W. L., Chandrasekhar, J., Madura, J. D., Impey, R. W. & Klein, M. L. Comparison of Simple Potential Functions for Simulating Liquid Water. J Chem Phys 79, 926–935, doi:10.1063/1.445869 (1983).

71 Huang, J. et al. CHARMM36m: an improved force field for folded and intrinsically disordered proteins. Nat Methods 14, 71–73, doi:10.1038/nmeth.4067 (2017).

72 Feller, S. E., Zhang, Y. H., Pastor, R. W. & Brooks, B. R. Constant-Pressure Molecular-Dynamics Simulation - the Langevin Piston Method. J Chem Phys 103, 4613–4621, doi:10.1063/1.470648 (1995).

73 Darden, T., York, D. & Pedersen, L. Particle Mesh Ewald - an N.Log(N) Method for Ewald Sums in Large Systems. J Chem Phys 98, 10089–10092, doi:10.1063/1.464397 (1993).

74 Phillips, J. C. et al. Scalable molecular dynamics with NAMD. J Comput Chem 26, 1781–1802, doi:10.1002/jcc.20289 (2005).

75 Shi, Y., Wang, L., Zhang, J., Zhai, Y. & Sun, F. Determining the target protein localization in 3D using the combination of FIB-SEM and APEX2. Biophys Rep 3, 92–99, doi:10.1007/s41048-017-0043-x (2017).

76 Tongren Yang et al. The microgravity enhanced polymer-mediated siRNA gene silence by improving cellular uptake. Biophys Rep, 6–0, doi:10.1007/s41048-020-00121-y (2020).

77 Scholz, B. A. et al. WNT signaling and AHCTF1 promote oncogenic MYC expression through super-enhancer-mediated gene gating. Nat Genet 51, 1723–1731, doi:10.1038/s41588-019-0535-3 (2019).

