## Supplemental Data for "Determining the architecture of nuclear ring of *Xenopus laevis* nuclear pore complex using integrated approaches"

### Supplementary Information

#### Supplementary Figures

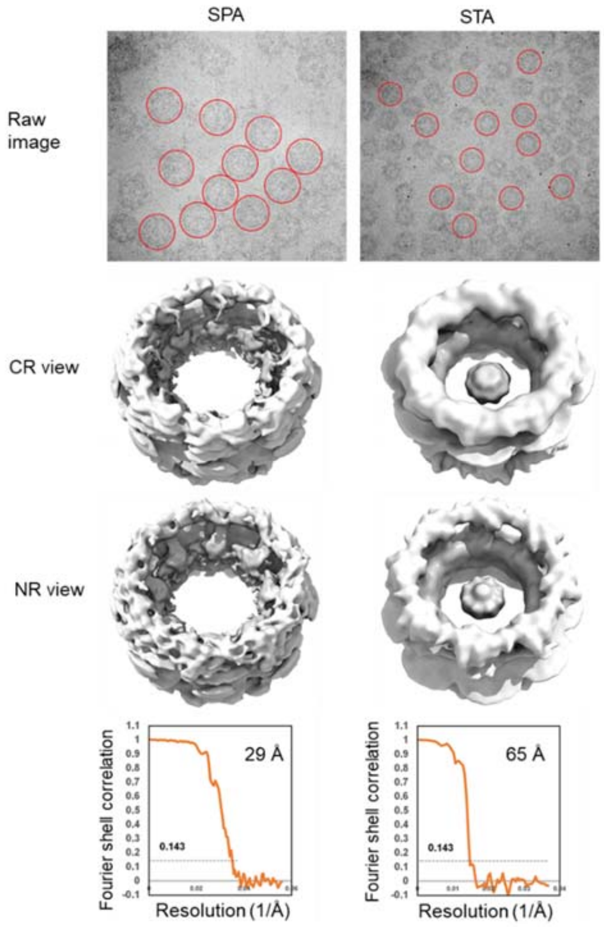


**Fig. S1 Comparison of *X. laevis* NPCs solved by single particle analysis (SPA) and sub-tomogram averaging (STA).** Raw images of SPA and STA samples have been low pass filtered to 40 and 50 Å respectively for better visualization, and representative NPC particles were marked by red circles. *X. laevis* NPCs reconstructed using the SPA and STA strategies were shown in both the CR view and NR view, indicating high similarity between the two maps. The gold standard FSC (Fourier Shell Correlation) curves are shown below with the 0.143 resolution-criteria indicated.

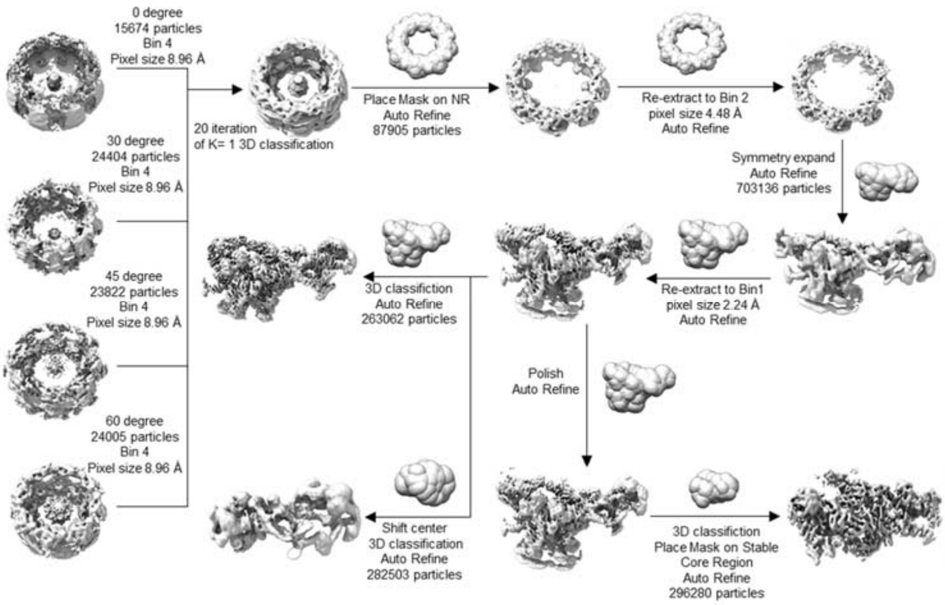


**Fig. S2 Data processing workflow of cryo-EM SPA in this work.**

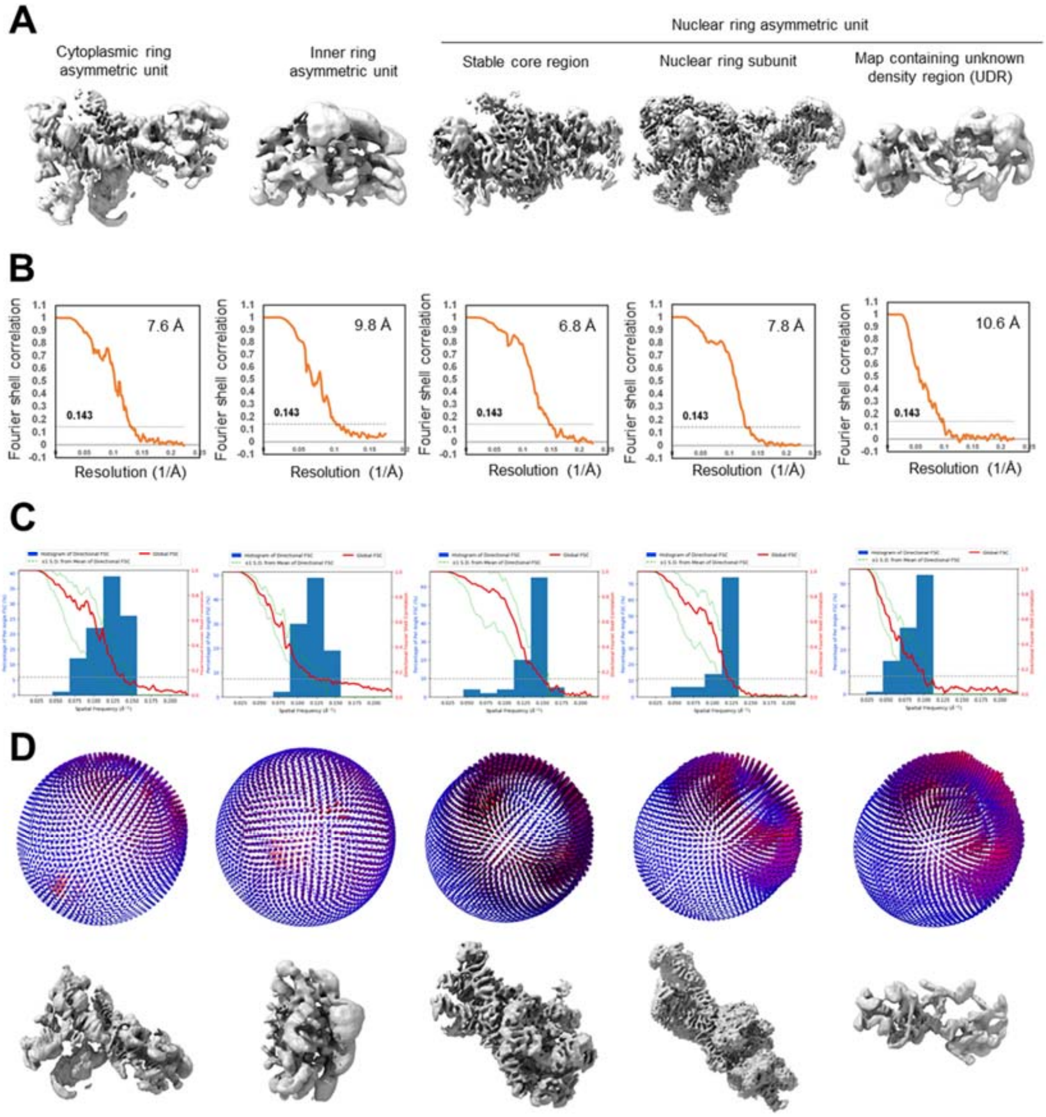


**Fig. S3 Overall qualities of cryo-EM maps in different regions of *X. laevis* oocyte NPC.** **(A)** Different regions of the NPC were refined using localized masks, with resolution ranging between 6.8 Å and 10.6 Å. (**B-D**) The gold standard FSC (Fourier Shell Correlation) with the 0.143 resolution-criteria, 3DFSC estimates, and angular distributions for each map were shown.

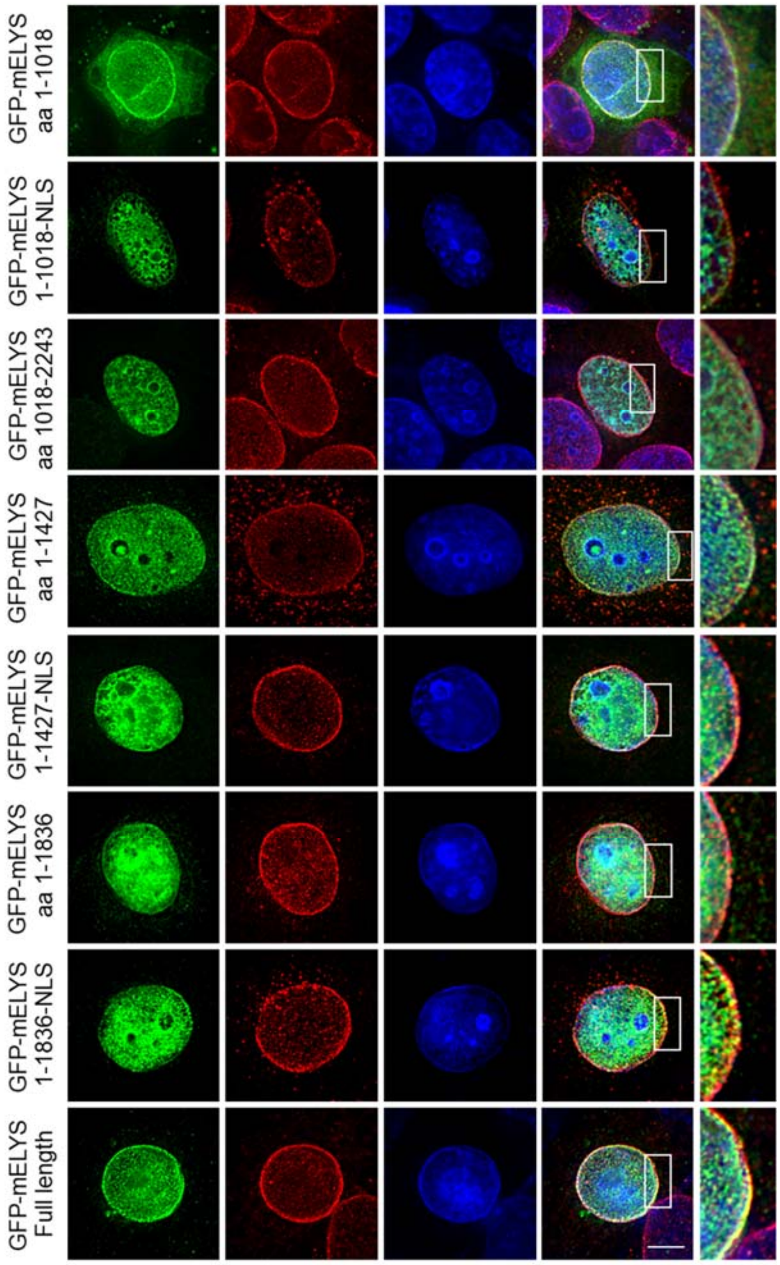


**Fig. S4 Localization of full-length and truncated mutants of ELYS in cells.** Full-length and truncated mutants (with or without NLS) of GFP-tagged mouse ELYS (mELYS) were expressed in HeLa cells for 24 h, followed by fixation with methanol and immunostaining with the anti-NPC antibody mAb414 (red). DNA was count-stained with DAPI. Scale bar, 10 μm.

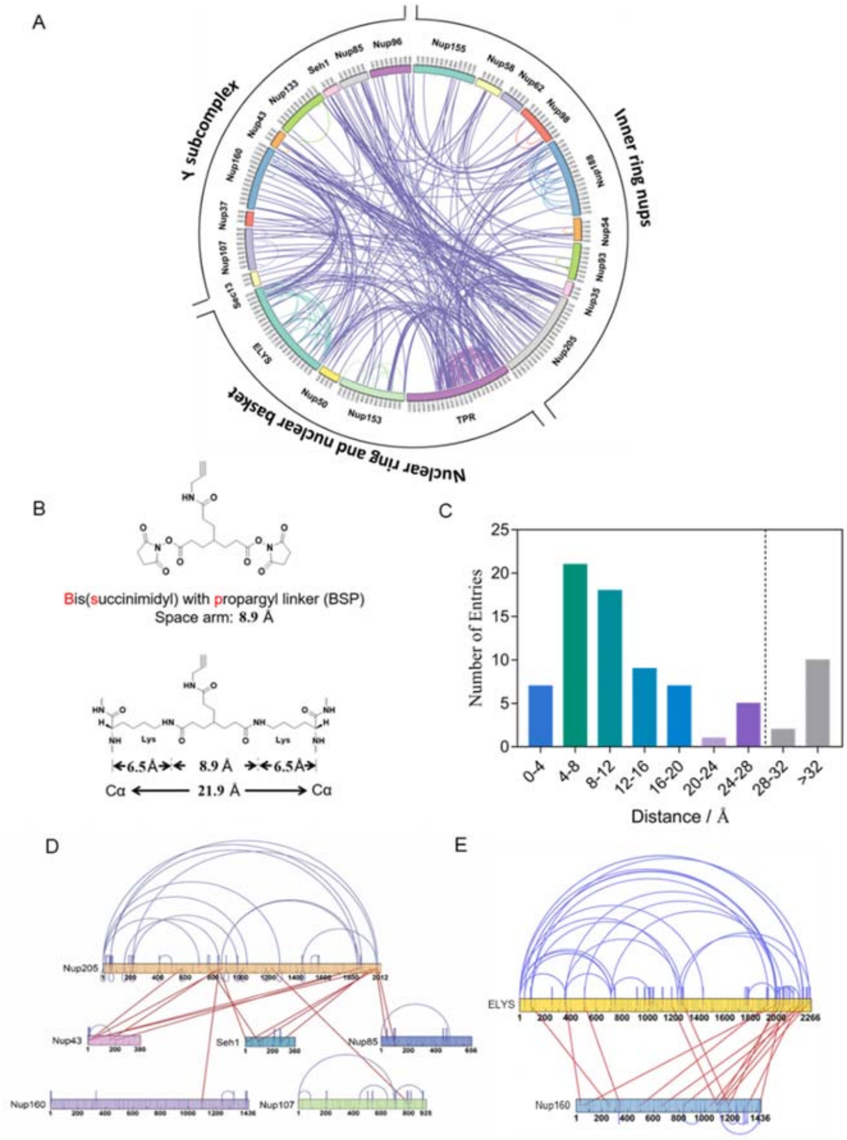


**Fig. S5 Cross-linking of NR components by *in situ* chemical cross-linking and mass spectrometry. (A)** The connectivity of NR proteins identified by chemical cross-linking and mass spectrometry. (**B**) The chemical structure and crosslinking restraint of BSP cross-linker. (**C**) The distance distribution of all cross-links that could be mapped to the known high-resolution structures of Nups (PDB entry:3TJ3, 4LIR, 5A9Q, 5IJN, 5IJO, 5TO5). **(D)** Schematic view of cross-linking sites of Nup205 with other Nups revealed by CX-MS. (**E**) Cross-linking mass spectrometry shows that ELYS binds with Nup160 at multiple sites.

### Supplementary Tables

**Table S1 Nomenclature for the Nups of CR, Y complex, NR and basket, in *Saccharomyces cerevisiae*, *Schizosaccharomyces pombe*, *Homo sapiens* and *Xenopus laevis*.**

| | *S. cerevisiae* | *S. pombe* | *H. Sapiens* and *X. laevis* |
|---|---|---|---|
| | — | — | Nup358 |
| | Nup159 | Nup146 | Nup214 |
| Cytoplasmic Ring (CR) | Nup82 | Nup82 | Nup88 |
| | Nsp1 | Nsp1 | Nup62 |
| | Nup192 | Nup186 | Nup205 |
| | Nup188 | Nup184 | Nup188 |
| | Nup85 | Nup85 | Nup85 |
| | Seh1 | Seh1 | Seh1 |
| | Nup120 | Nup120 | Nup160 |
| | Sec13 | Sec13 | Sec13 |
| Y complex | Nup145C | Nup189 | Nup96 |
| | Nup84 | Nup107 | Nup107 |
| | Nup133 | Nup131 | Nup133 |
| | — | — | Nup43 |
| | — | Nup37 | Nup37 |
| | — | ELY5 | ELYS |
| | Nup192 | Nup186 | Nup205 |
| Nuclear Ring (NR) and basket | Nup1 | Nup124 | Nup153 |
| | Nup2 | Nup61 | Nup50 |
| | Mlp1<br>Mlp2 | Nup211 | TPR |

**Table S2 Statistics of cryo-SPA data collection and image processing.**

| | | | | | |
|---|---|---|---|---|---|
| **Data acquisition** | | | | | |
| Microscope | Titan Krios G2 | | | | |
| Voltage (kV) | 300 | | | | |
| Detector | Gatan K2 | | | | |
| Energy filter | Gatan GIF Quantum, 20 eV | | | | |
| Mode | Super resolution | | | | |
| Pixel size (Å) | 2.24 | | | | |
| Stage tilting angle | 0°/ 30°/ 45°/ 60° | | | | |
| Exposure per tilt (e/Å²) | 60 / 60/ 80/ 100 | | | | |
| Number of images | 5971 | | | | |
| Defocus range (μm) | -1 ~ -4 | | | | |
| Software | SerialEM | | | | |
| **Reconstruction** | | | | | |
| Software | RELION-3.0 | | | | |
| Data set | NR stable core region | NR subunit | map containing UDR | CR asymmetric unit | IR asymmetric unit |
| Final number of particles | 296280 | 263062 | 282503 | 521065 | 703128 |
| Symmetry | C1 | C1 | C1 | C1 | C1 |
| Final resolution (Å) | 6.8 | 7.8 | 10.6 | 7.6 | 9.8 |
| Map pixel size (Å) | 2.24 | 2.24 | 2.24 | 2.24 | 2.24 |
| Map sharpening B-factor (Å²) | -328 | -482 | -1095 | -352 | -264 |

**Table S3 Statistics of cryo-ET data collection and image processing.**

| | |
|---|---|
| **Data acquisition** | |
| Microscope | Titan Krios G2 |
| Voltage (kV) | 300 |
| Detector | Gatan K2 |
| Energy filter | Gatan GIF Quantum, 40 eV |
| Mode | Counting |
| Pixel size (Å) | 3.4 |
| Tilt step & range | -60° ~ +60°, 3° step, start at 0° |
| Tilt scheme | Dose-symmetric |
| Number of tilt-series | 198 |
| Exposure per tilt (e/Å²) | 3.5 |
| Defocus range (μm) | -1.5 ~ -3 |
| Software | SerialEM |
| **Reconstruction** | |
| Software | Dynamo v1.1.509 |
| | RELION v3.0 |
| | Warp 1.0.7 & 1.0.9 |
| Data set | *X. laevis* NPC |
| Final number of particles | 1360 |
| Symmetry | C8 |
| Final resolution (Å) | 65 |
| Map pixel size (Å) | 13.6 |

### Supplementary Movies

**Movie S1 Overview of reconstruction of *X. laevis* NPC cryo-EM map.**

### Supplementary Script

**Script S1 The modified block-based reconstruction script for symmetry expanding of NR asymmetric unit.**
